# Monitoring Functional Post-Translational Modifications Using a Data-Driven Proteome Informatic Pipeline

**DOI:** 10.1101/2022.11.09.515610

**Authors:** Payman Nickchi, Uladzislau Vadadokhau, Mehdi Mirzaie, Marc Baumann, Amir A. Saei, Mohieddin Jafari

**Author notes:** Correspondence to Amir Ata Saei or Mohieddin Jafari. These authors contributed as co-last authors (Amir A. Saei, Mohieddin Jafari).

## Abstract

Post-translational modifications (PTMs) are of significant interest in molecular biomedicine due to their crucial role in signal transduction across various cellular and organismal processes. Characterizing PTMs, distinguishing between functional and inert modifications, quantifying their occupancies, and understanding PTM crosstalk are challenging tasks in any biosystem. Studying each PTM often requires a specific, labor-intensive experimental design. Here, we present a PTM-centric proteome informatic pipeline for predicting relevant PTMs in mass spectrometry-based proteomics data without prior information. Once predicted, these in silico identified PTMs can be incorporated into a refined database search and compared to measured data. As a practical application, we demonstrate how this pipeline can be used to study glycoproteomics in oral squamous cell carcinoma based on the proteome profile of primary tumors. Subsequently, we experimentally identified cellular proteins that are differentially expressed in cells treated with multikinase inhibitors dasatinib and staurosporine using mass spectrometry-based proteomics. Computational enrichment analysis was then employed to determine the potential PTMs of differentially expressed proteins induced by both drugs. Finally, we conducted an additional round of database search with the predicted PTMs. Our pipeline successfully analyzed the enriched PTMs, and detected proteins not identified in the initial search. Our findings support the effectiveness of PTM-centric searching of MS data in proteomics based on computational enrichment analysis, and we propose integrating this approach into future proteomics search engines.

## Main

Proteins are the primary functional units of cellular systems, but they often become active only after post-translational modifications. In addition to regulation of protein activity, their function, stability/solubility, interactions with other biomolecules and their cellular localization are governed by transient modulation of post-translational modifications (PTMs)^1, 2^. By regulating such diverse characteristics, PTMs can modulate the involvement of proteins in biochemical reactions, signaling, transport, structural remodeling, gene regulation, cell motility and cell death^3, 4^. Due to the importance of PTMs in signal transduction in health and disease, elucidating the mechanisms and the kinetics of PTMs have turned into an active research area^5–9^.

Analysis of PTMs would provide valuable information regarding the status and function of proteins upon diverse perturbations^10^. Therefore, understanding the nature, quantity, and temporal progression of PTMs has arguably been one of the most substantial contributions of MS-based proteomics to modern biology^1^. However, the sub-stoichiometric nature and dynamic regulation of PTMs makes it challenging to capture and detect PTMs^11^. Thus, unique enrichment techniques and sample-processing workflows are often required for enriching PTMs before analysis by mass spectrometry^12^.

Experimental techniques exploit the unique chemical properties of a given PTM for their enrichment. For example, at both protein and peptide levels, PTM-directed antibodies can be used to enrich a specific chemical group within a given proteome^1^. Another routinely used strategy involves the enrichment of phosphorylated peptides (and/or proteins) using metal oxide resins, such as titanium and zirconium^13^. These enrichment strategies are particularly effective when modulation of a certain PTM is expected; for example, when investigating the function of a kinase, modulation of phosphorylation levels is an expected outcome.

The incorporation of known modification sites encoded in formats such as PEFF presents a promising avenue for refining and optimizing PTM searches^14^. However, it is essential to consider the potential challenges associated with enlarging the search space, as the inclusion of more PTMs in database searches can lead to increased time constraints and strain on search engine^15^. It should also be noted that including any extra PTMs increases the chance of false identifications in a given database search and thus increasing the burden of proof for PTM identification. Therefore, only most common PTMs, such as asparagine deamidation^16^, methionine oxidation and cysteine carbamidomethylation are usually included in routine database searches.

Altogether, due to a lack of prior knowledge on the most important PTMs in a particular study condition, many PTMs are usually not monitored. Although no practical issues exist in the biochemical characterization of stable and common PTMs such as phosphorylation or acetylation, researchers do not monitor them without prior knowledge. Despite the presence of proteome-wide PTM approaches such as ModifiComb^17, 18^, the analysis of less-common or unstable PTMs still remains challenging and needs a more complex study design, especially for PTMs without highly specific antibodies or reagents.

Here, we present a thorough analysis of the advancements introduced by the PEIMAN2 R package in the realm of PTM-centric discovery proteomics. We demonstrate the effectiveness of the software package by conducting two extensive case studies: one utilizing external public data and the other leveraging in-house data. These case studies collectively serve as strong evidence, validating the utility and potential applicability of our proposed pipeline in practical proteomic research. The first case study highlights the predictability of PEIMAN2 in glycoproteome profiling, demonstrated by its application to the analysis of oral squamous cell carcinoma (OSCC), utilizing pre-existing proteomic and glycoproteomic data^19^. Subsequently, an in-house case study focuses on the identification of mechanistic proteins subject to differential expression by the multikinase inhibitors dasatinib and staurosporine. This investigation involves deep expression profiling of the A549 cell line, serving as a representative model for lung cancer. In both studies, a PEIMAN2-based search pipeline (as illustrated in **Fig. 1**) is employed to identify the enriched PTMs among the differentially expressed proteins. Finally, these enriched PTMs are seamlessly integrated into a refined database search, enabling an evaluation of the PEIMAN2 pipeline in the domain of PTM-centric discovery proteomics.

## Results

### PTM data extraction for PEIMAN2 R package

The PEIMAN2 R package contains as of September 2023 a database that encompasses 134,783 proteins from 14,457organisms, covering 515 PTM types. We compared the distribution of the 10 most prevalent PTMs across all known organisms to gain insight into the prevalence of PTMs in the current version of UniProtKB/Swiss-ProtKB/Swiss-Prot vs 2015 version. The ten most common PTMs in the current version of UniProtKB/Swiss-ProtKB/Swiss-Prot are “Phosphoprotein”, “Disulfide bond”, “Phosphoserine”, “Glycoprotein”, “Acetylation”, “Phosphothreonine”, “Ubl conjugation”, “Lipoprotein”, “N6-acetyllysine”, and “Methylation” (**Fig. 2A**). The term “Phosphoprotein” had a frequency of 48,934 occurrences which is over 5 times greater than the “Methylation” term, which ranked as the tenth most frequent PTM with a frequency of 8,589 occurrences and displayed the highest increase rate (1.5-fold) over the past decade. The growth trends of these ten most common PTMs in the current UniProtKB/Swiss- ProtKB/Swiss-Prot database compared to previous versions exhibit variations in their rates (**Fig. 2B**). Over the course of the past seven years, we expected to discover a consistent pattern of growth among the top 10 selected PTMs. However, the rates of growth are not consistent among all terms. For example, “Methylation”, “Phosphothreonine”, and “Phosphoserine” terms have a higher rate of growth compared to the other PTM-terms. This difference in the rate of growth might be related to two reasons. First, the identification of some PTMs is subjected to experimental limitations, therefore we cannot expect a consistent rate of growth. Second, the assigned biological activities of some PTMs such as phosphorylation has attracted more research attention than some other PTMs.

**Figure 2:**
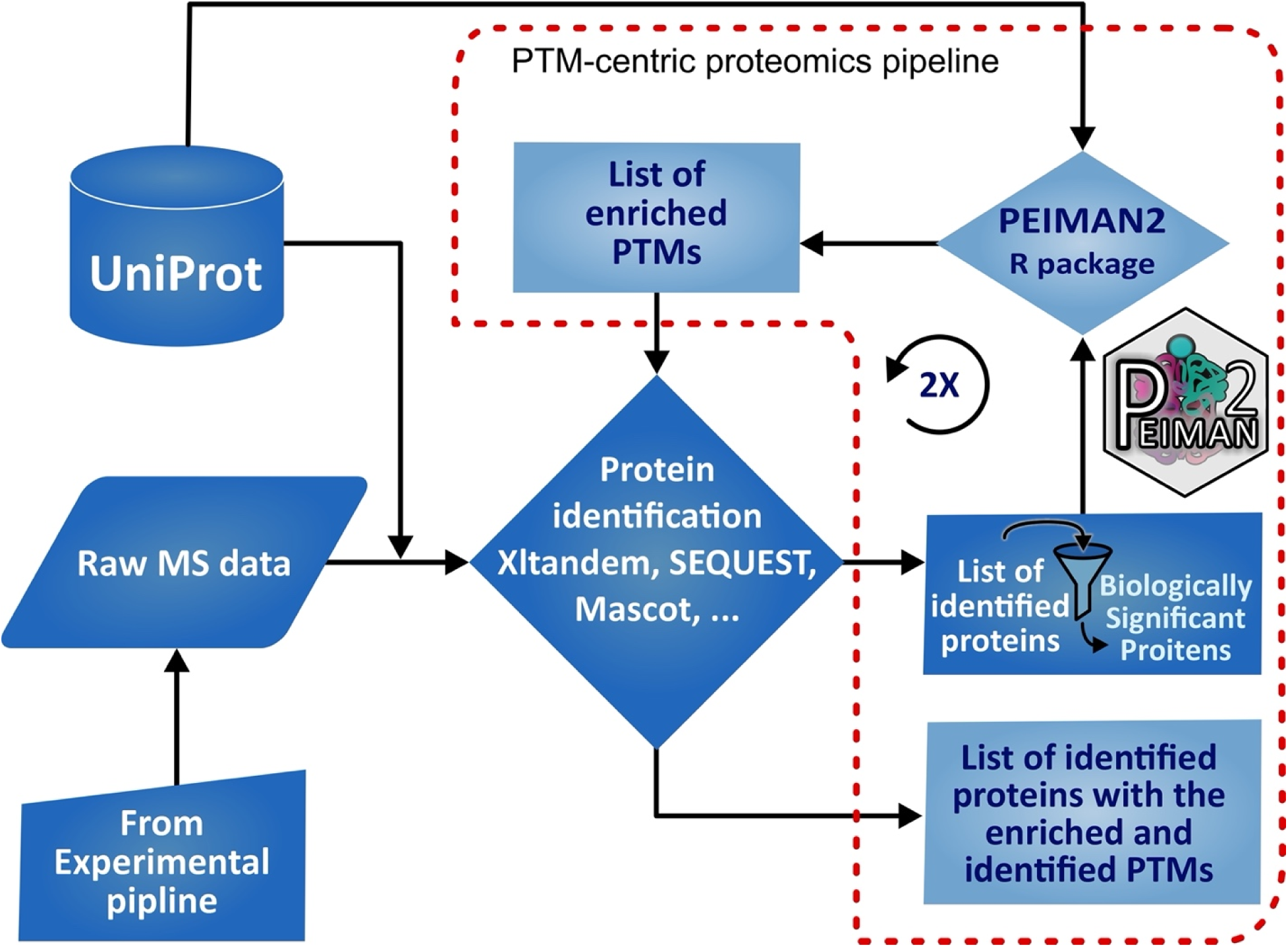
An informatic pipeline for PTM-centric proteomics using PEIMAN2 R package. The red dash line area delineates the inputs and outputs of PEIMAN2 R package, forming a PTM-centric proteomics pipeline.

Furthermore, because PEIMAN2 relies on annotated PTMs in the UniProtKB/Swiss-Prot database, we conducted an analysis of PTM distribution across various species within UniProtKB/Swiss-Prot. Tree maps were employed to visually represent this distribution across eight model organisms, each selected to represent diverse taxonomic branches (Supplementary Figure 1 in Supplementary data 5). The observed PTM distribution among these distinct model organisms suggests the potential utility of PTMs for taxonomic discrimination within the broader context of the tree of life, thereby prompting further avenues of research. Additionally, a t-distributed Stochastic Neighbor Embedding (t-SNE) plot^20^ was generated to provide a comprehensive visualization of how the PTM profiles of species enable the classification of these organisms into four major super kingdoms (Supplementary Figure 2 in Supplementary data 5).

### PEIMAN2 R package functionality

Previously, we introduced a computational enrichment analysis for PTMs by a standalone software called “Post-translational modification Enrichment Integration and Matching Analysis” or PEIMAN, to facilitate singular enrichment analysis (SEA) based on PTMs in proteomics studies^21^. SEA^22^ is a popular method providing insight into biological pathways altered in disease or under various perturbations. The idea of SEA is to check whether the genes/proteins with a specific biological feature in a given list are occurring more frequently than by pure chance. As simple and powerful as this approach is, there are some known drawbacks to it, including the difficulty of identifying significant signals from noise, subjective interpretations among biologists, and for the same data getting different final list of significant genes/proteins among different laboratories. Gene set enrichment analysis (GSEA) offers an alternative solution to these challenges. Therefore, we developed a new enrichment method specific to PTMs in proteins, called Protein Set Enrichment Analysis (PSEA), and made it available as an R package (see Methods section). This tool is now accessible to a broader community of researchers.

The PEIMAN2 R package offers a wider range of features and functionalities compared with the PEIMAN standalone software. First note that SEA related functions are still included in the PEIMAN2 R package and can be utilized by calling `runEnrichment()` and `plotEnrichment()` function. As a new feature, PEIMAN2 package implements PSEA as an additional tool for proteomics studies based on PTMs. `runPSEA()` function in the package allows the user to perform a PSEA analysis on a given list of proteins (e.g., UniProt IDs) for a specific organism. This function requires a list of protein accession codes along with taxonomy name of the organism. Some additional parameters of the function include enrichment weighting (refer to the methods section for more details), number of permutations to estimate the false discovery rate, FDR (default number of permutations is 1000), choice of the method to adjust p-values, and a controlling cut off to include specific PTMs with a certain occurrence rate in the analysis. A table of enriched proteins along with their enrichment and normalized enriched score, adjusted p-value, FDR, and proteins in the leading-edge is produced. For each PTM, the leading-edge proteins are the proteins that show up in the ranked list at /or before the point where the enrichment score (ES) reaches its maximum deviation from zero.

There are two functions available in the package to visualize the results of PSEA, `plotPSEA()` and `plotRunningScore()`. The former is employed for generating plots depicting the outcomes of a single PSEA analysis or for merging the findings from two separate PSEA analyses. These plots effectively display the normalized enrichment scores associated with each PTM. The latter function is designed to produce running enrichment score plots for each PTM featured in the table generated by the `runPSEA()` function. In each plot, x-axis is the sorted protein list and y-axis is the enrichment score. The leading-edge proteins are shown with a rug plot on the x-axis (for example see Supplementary Figure 3 and 4 in Supplementary data).

In PTM-centric proteomics, we recommend integrating PEIMAN2 results into the workflow of mass spectrometry data analysis. To do this, one can utilize SEA or PSEA to generate a list of enriched terms. These are PTM terms of which the counts in a given list of protein is statistically significant. The SEA and PSEA are two methods to obtain the list of enriched PTM terms (see methods section for more details). The enriched terms can then be used to extract the subset of protein modifications. These modifications can be used to search in a proteomics search engine software to gain more insight into designing the experiment and investigating the effect of a treatment on PTMs. For this purpose, we included functions to prepare results for such a research in MaxQuant software. The results of SEA or PSEA can be passed to `sea2mass()` or `psea2mass()` functions, respectively, to extract a subset of protein modifications. This subset of chemical modifications can be used to parametrize the search engine for mass spectrometry data, such as MaxQuant. For more information, we have provided a detailed vignette manual along with the package and a Readme page on PEIMAN2’s GitHub directory (https://github.com/jafarilab/PEIMAN2).

### Glycoproteomics case study with PEIMAN2

To evaluate and benchmark PEIMAN2’s efficiency in prediction of relevant PTMs in a given proteomics study, we applied PEIMAN2 to a recent study by Carnielli *et al.* ^19^, where they perform proteomics and glycoproteomics profiling on samples from oral squamous cell carcinoma (OSCC) patients. This study was based on the existing understanding of the significant role of glycosylation in regulating crucial factors such as altered adhesion behavior, migratory tendencies, and metastatic advancement of oral cancer cells. In this study, primary tumor tissues, extracted through surgical procedures from OSCC patients including those with and without lymph node metastasis, were subjected to proteomics and glycoproteomics analyses. Through clustering analysis, the study quantitatively juxtaposed the N-glycome and N- glycoproteome data across diverse patient groups. These analyses, coupled with the exploration of an array of clinicopathological features using patient metadata, revealed substantial changes in the abundance of several N-glycopeptides, establishing a compelling connection between glycoproteins and patient survival outcomes.

To assess PEIMAN2’s ability in computationally enriching and predicting glycosylation within the glycoproteome profiling dataset, we applied PEIMAN2 to identify the enriched PTMs (**Fig. 3A** and **3B**). The analysis unveiled a pronounced enrichment of glycoprotein and GPI-anchor amidated alanine, asparagine, and cysteine – findings that aligned with the anticipated outcomes. Therefore, this analysis serves as a proof-of principle. Next, we extended our assessment to the routine proteome profiles of both patient groups to validate our approach in an experimental setting where no prior knowledge of involved PTMs, in this case, glycosylation, was available. Impressively, the PTMs associated with glycoproteins were enriched significantly as a major PTM among the proteins exhibiting differential expression in the typical proteome profile —without specific experimental enrichments targeting glycosylation (**Fig. 3C** and **3D**). This finding shows that if *a priori* information on the involvement of glycosylation on the progression of OSCC was missing, PEIMAN2 would have been capable of correctly identifying glycosylation as a major PTM solely by using the routine proteomics dataset, guiding the authors to perform subsequence glycoproteomics studies. This insight is valuable in analogous situations where prior knowledge about the significance of various PTMs in diverse biological and disease contexts is lacking. Importantly, the matched proteomics and glycoproteomics data in this study was beneficial and could be used as a perfect benchmark to showcase the applicability of PEIMAN2.

**Figure 3:**
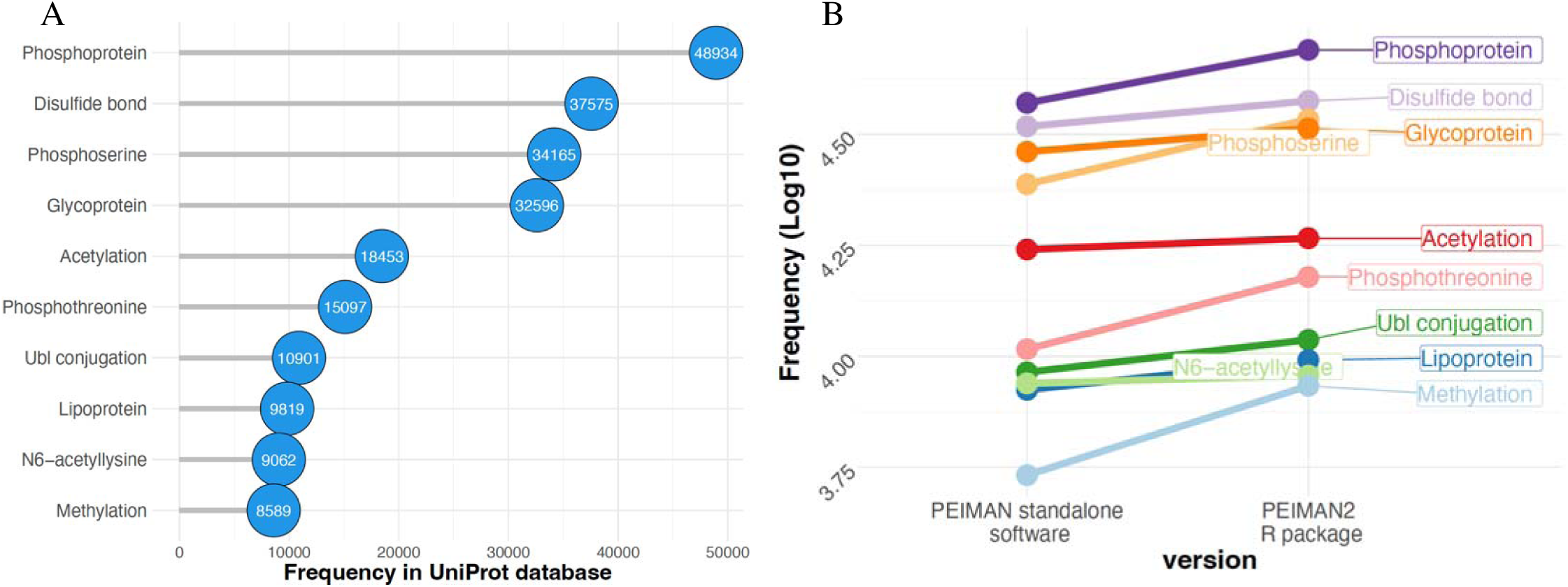
An update on PTM statistics based on UniProtKB/Swiss-Prot database. (A) The top ten most common PTMs in UniProtKB/Swiss-Prot across all available species. (B) The frequency changes of the ten most common PTMs in PEIMAN standalone software (2015) vs PEIMAN2 R package (2023). Note that the database version is denoted on the x axis and the y axis represents the frequency of PTMs. To better highlight the changes, the frequency of PTMs is shown in log-10 base.

Subsequently, we continued with an in-house investigation on deep expression profiling of cells treated with two kinase inhibitors that inhibit protein phosphorylation. The overarching goal was to predict the modifications in the expression proteomics data and to pinpoint mechanistic proteins that exhibit differential expression in response to treatment, reflecting potential pathways and mechanisms influenced by drug interventions.

### Multikinase inhibitors case study with PEIMAN2

Since both drugs are multikinase inhibitors, the treatment would be expected to modulate some phosphorylation events and/or modification occupancies, as shown before for AKT1/2 inhibitor and ipatasertib^2^. Therefore, we expected to see an enrichment of phosphorylation PTM terms after applying PEIMAN2 on the differentially expressed proteins in response to both drugs. We tested four increasing concentrations of both drugs in an A549 cell line model of lung cancer taking advantage of TMTpro 16 multiplexing^23^. As expected, a higher number of differentially expressed proteins were identified at higher concentrations of drug. Then, we tested PEIMAN2 on all concentrations of drugs and checked the number of differentially expressed proteins with annotated PTM term changes at different drug concentration levels based on enrichment analysis. <u>The plots for dasatinib and staurosporine drugs (</u>**Fig. 4A** and **4B**<u>) illustrate</u> <u>the presence of various PTMs on differentially abundant proteins when comparing the highest</u> <u>concentration of drugs (Conc.4) versus control. As drug concentration increases from Conc.1 to</u> <u>Conc.4, both the number of differentially expressed proteins (</u>**Fig. 4C** and **4D**<u>) and potential</u> <u>protein targets with specific PTM terms (</u>**Fig. 4E** and **4F**<u>) show an upward trend.</u> Therefore, we considered Conc.4 for the downstream analyses including a refined MaxQuant database search. For emphasis, we tried to detect actual PTMs in mass spectrometry data based on the PTMs that were enriched in the identified proteins using PEIMAN2.

**Figure 4:**
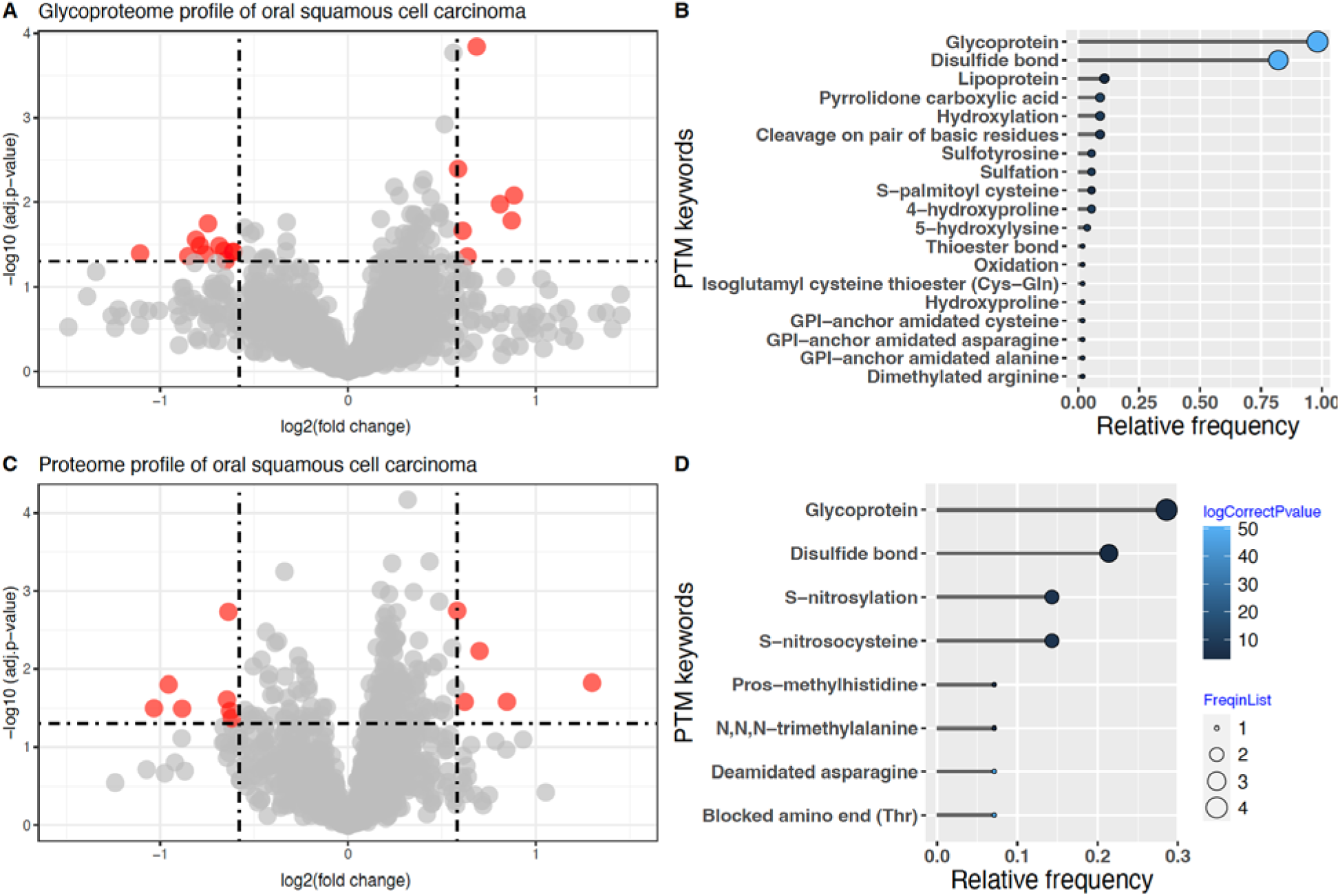
Proteome and glycoproteome profiling in OSCC. (A) Volcano plots illustrate the OSCC glycoproteome, and (C) the proteome, with data points color-coded to indicate differentially expressed proteins. Enriched PTMs in the (B) glycoproteome and (D) proteome profiles are presented using PEIMAN2 with lollipop plots. The p-values are represented through a gradient from black to light blue, as shown by the spectral color bar. The size of each circle in the lollipop plot corresponds to the number of proteins with a corresponding PTM in our protein list.

More specifically, for obtaining the most probable modifications changing under the treatment, at the first step of analysis, we implemented PSEA method by calling `runPSEA()` function on the samples treated with the highest concentration of drugs, to identify enriched modification terms. As for permutation, we considered randomly permuting scores of proteins 1000 times to adjust for FDR. A significance level of 5 percent along with a reasonable cut-off for PTM frequency in UniProtKB/Swiss-Prot was applied to each drug list, separately. The exact modification of each enriched PTM was obtained by calling `psea2mass()` function. The top five modifications for dasatinib were: ‘O-phospho-L-threonine’, ‘N6-acetyl-L-lysine’, ‘N-acetyl-L- alanine’, ‘O4-phospho-L-tyrosine’, and ‘N-acetyl-L-methionine’. The corresponding PTMs are: ‘Phosphothreonine’, ‘N6-acetyllysine’, ‘N-acetylalanine’, ‘Phosphotyrosine’, ‘N- acetylmethionine’. On the other hand, the top five modifications for staurosporine were: ‘O- phospho-L-serine’, ‘O-phospho-L-threonine’, ‘O4-phospho-L-tyrosine’, ‘N-acetyl-L-alanine’, and ‘N-acetyl-L-methionine’. The corresponding PTMs are: ‘Phosphoserine’, ‘Phosphoprotein’, ‘Acetylation’, ‘Phosphothreonine’, and ‘N6-acetyllysine’. In Supplementary data 5, Supplementary Figure 3 and 4, we show the running score plot of the top five modifications shown in the integrated normalized enrichment score plot for dasatinib and staurosporine, respectively. In summary, PSEA obtained from 1000 permutations shows enriched phosphorylation for both drug treatments (P-value < 0.05), however, the top PTMs for these kinase inhibitors were distinct and specific (**Fig. 4G**). In the next step, we performed a refined search using MaxQuant, where the top 5 modifications suggested by PEIMAN2 were added as further variable modifications (other parameters were kept constant as with the initial search).

### Number of proteins, peptides, and sequence coverage before/after using PEIMAN2

First, we performed a quality control to ensure that the number of unique peptides, sequence coverage, and q-values are not significantly changed after including additional enriched modifications suggested by PEIMAN2 on the proteomics search engine software, i.e., MaxQuant. Note that the re-searching parameters (except the selected modifications) as well as filtering criteria of selecting the identified proteins remained the same both before/after including PEIMAN2 suggestions. Figure 5 shows the box plot of sequence coverage of proteins for each unique number of peptides colored by their q-value before/after including additional enriched modifications suggested by PEIMAN2. The results of analysis before/after applying PEIMAN2 for dasatinib and staurosporine are presented in panels (A-B) and (C-D), respectively. For both drugs, the median of sequence coverage at each unique number of peptides was similar before and after applying PEIMAN2 suggested modifications.

**Figure 5:**
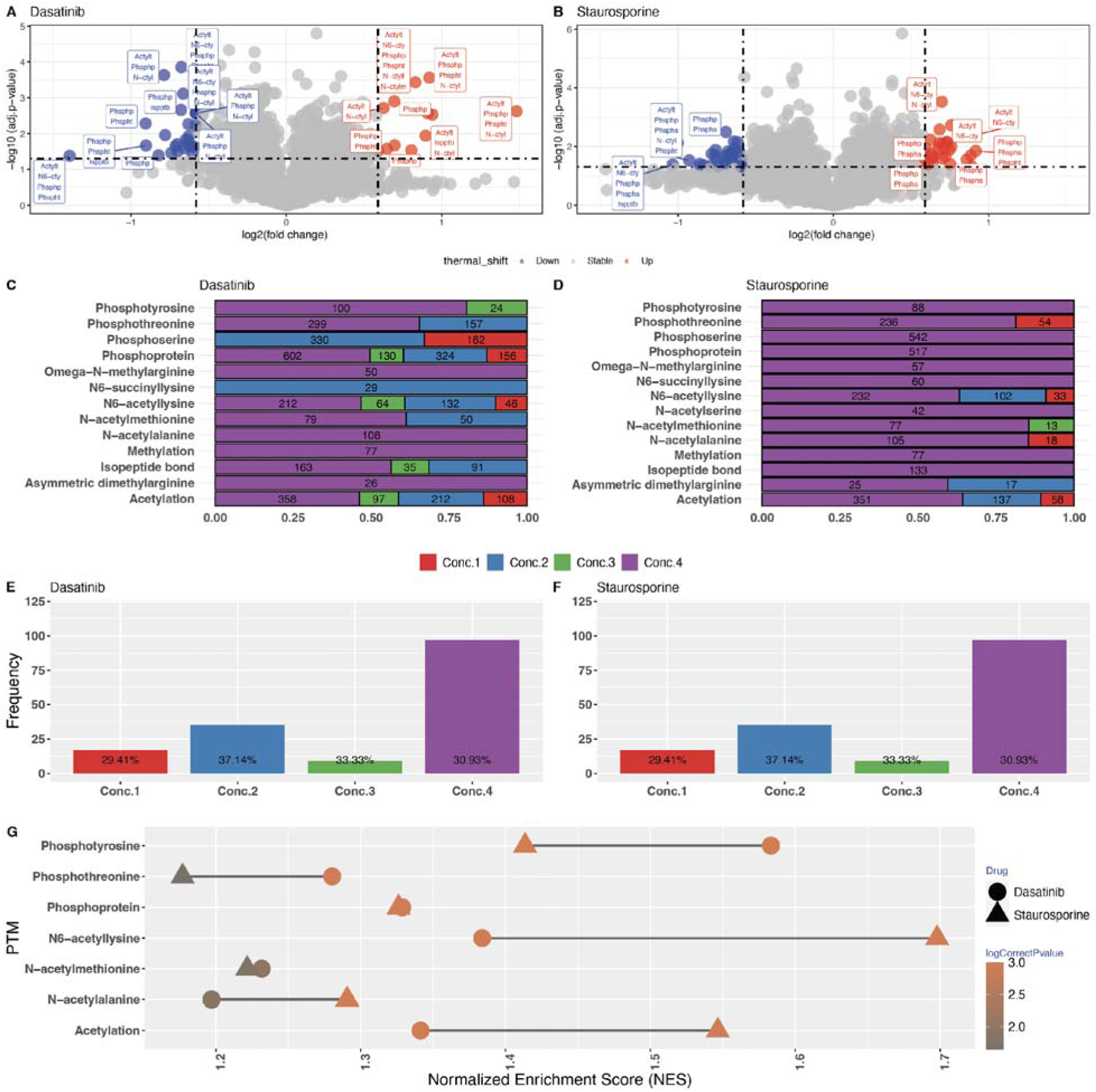
Volcano plot of fold change versus p-value colored by the sign of protein expression for both drugs; the PTMs annotated in UniProtKB/Swiss-Prot for each protein are shown in rectangles; (A) Dasatinib, (B) Staurosporine. The results are shown for concentration 4 vs. vehicle treated cells. Bar plot of number of differentially expressed proteins with PTM at four concentration levels of both drugs (see Supplementary data 1 and 2 for more details); (C) Dasatinib, (D) Staurosporine. The value of absolute numbers of differentially expressed proteins carrying different PTMs at each concentration level is labeled in the bars of the plot; (E) Dasatinib, (F) Staurosporine. The percentage of proteins with phosphorylation related PTMs are labeled on the bars. (G) Integrated normalized enrichment score (NES) plot for both drugs colored by corrected p-value. The data points for dasatinib and staurosporine are plotted with a filled circle and triangle, respectively. The points are colored with their corrected p-value presented in log10 scale.

Then, we performed another quality control to check the number of proteins that were identified before/after using PEIMAN2 and checked whether the newly identified proteins carry any PTMs (**Fig. 5E** and **5G**). For dasatinib 126 new proteins were identified by including the modifications suggested by PEIMAN2. On the other hand, 52 proteins that were previously identified were lost in the refined search, suggesting that their corresponding peptides did not pass the 1% FDR threshold set in MaxQuant search. The majority of disappeared proteins were identified with two peptides in the initial search, with a median score value of 2.861 and small Q values. Note that 86% of proteins after including the suggested modification in the search had more than two peptides. There were 6,418 proteins in common before/after using PEIMAN2. For staurosporine, refined searched yielded 5,903 proteins that were also found in the initial database search. Furthermore, 123 proteins were identified that were not included in the initial search results, and 28 proteins were not found in the refined search after including PEIMAN2 suggested modifications. When comparing the results of the experiment before/after including PEIMAN2 suggestions, this result implies that more than 97 percent of the proteins that were quantified before/after PEIMAN2 implementation in the refined database search are the same. However, as a result of employing PEIMAN2, we now have information regarding the PTM status of previously identified proteins in addition to previously undiscovered proteins. It was interesting to investigate the types and frequency of modifications in the newly identified proteins after searching MaxQuant by considering PEIMAN2 suggestions (**Fig. 5F** and **5H**).

### PTMs on protein targets of drugs

We investigated the differential protein expression analysis before/after considering PEIMAN2’s suggestions in the database search. Figure 6 presents the fold change of proteins for each drug compared to control group, before/after considering PEIMAN2’s suggestions separately. The fold change is calculated as the log2 base of proportion of two sample means (Conc.4 vs control). The data points are colored depending on the results of a two-sample t-test (with equal variance assumption) before/after incorporating PEIMAN2’s suggestions in the database search. For example, purple color indicates that a protein was significantly different in Conc.4 group vs control group, both before/after using modification suggestions. These are drug- protein targets that were found repeatedly by MaxQuant and whose PTM status was clarified by PEIMAN2. The panels in Figure 6 shows the distinct modifications of proteins, if any.

**Figure 6:**
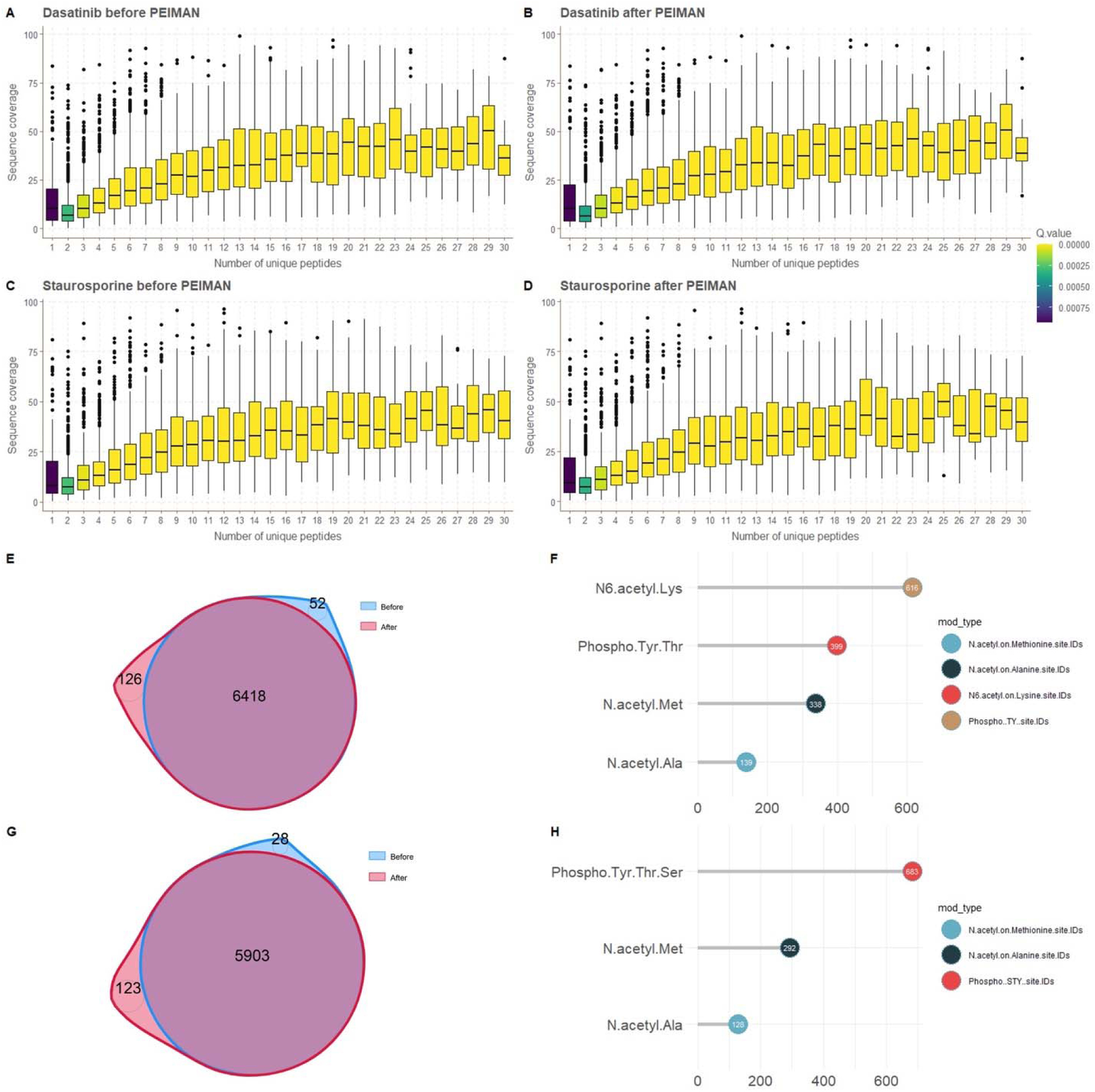
PEIMAN2 analysis-based before/after box plots for dasatinib and staurosporine. In these two pair plots (A:B and C:D), the effect of including additional enriched modifications suggested by PEIMAN2 on the search database by MaxQuant is depicted based on sequence coverage versus the number of unique peptides for both drugs separately. PEIMAN2 analysis- based before-and-after Venn diagrams and PTM frequency bar plots for dasatinib and staurosporine. The Venn diagrams (E) and (G), respectively, depict the number of proteins identified by MaxQuant for dasatinib and staurosporine before/after PEIMAN2 analysis. Based on PEIMAN2 analysis and re-searching the database by MaxQuant, the frequency plot of identified modifications of proteins is depicted for each drug (F and H).

**Figure 7:**
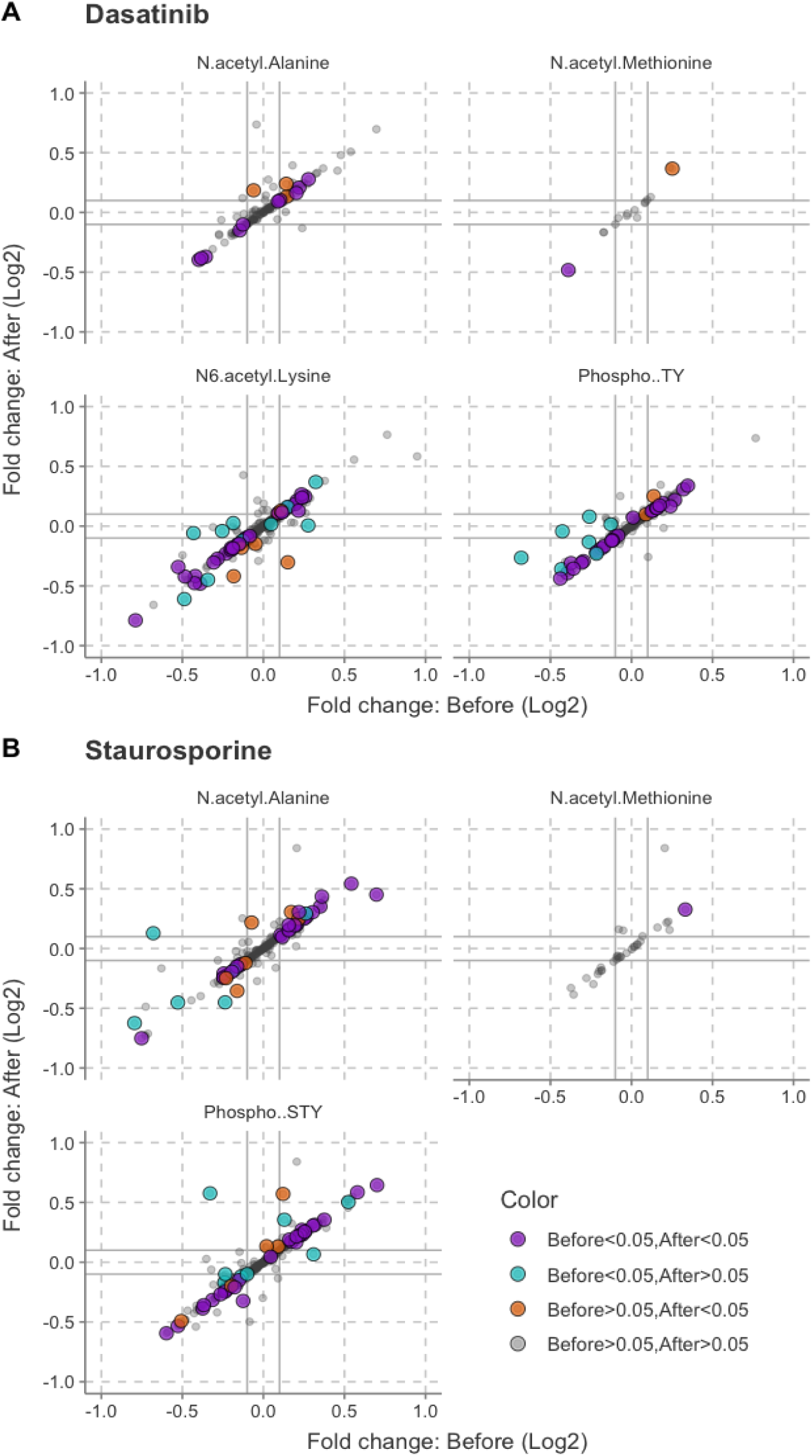
Differential analysis of proteins in response to the treatments before/after applying PEIMAN2. The fold changes of proteins with respect to untreated cells for each drug compared is displayed before/after considering PEIMAN2’s recommendations for incorporating enriched modifications in the database search. Four distinct colors were also used to depict the p-values of the t-statistics before/after PEIMAN2 analysis. Using shape icons, differential expressed proteins with distinct modifications are also indicated.

It is interesting to note that proteins that changed significantly before incorporating PEIMAN2’s suggestions are still statistically significant after considering the suggestions (purple color dots) with an enriched modification. In addition, the proteins that were not significantly changed before and became significant after including modification in MaxQuant search, are modified too. In comparison to other PTMs, the level of phosphorylated proteins is also more decreased or perturbed, necessitating further research to determine whether this phenomenon is because of the primary or secondary effects of inhibitors on protein targets. The above findings are consistent for both drugs. These results suggest that studies on drug targets and mechanism of action considering PTMs are helpful for identifying new proteins that are involved in drug mechanism of action. Basically, by including relevant PTMs in database search, the results of studies on different perturbations can be brought closer to reality using PEIMAN2.

### Monitoring of PTM changes after PEIMAN2 implementation

Finally, we investigated the perturbation of the quantified PTMs upon treatment with dasatinib and staurosporine at the peptide level. First, the intensity of a given peptide was normalized by total intensity for each TMT channel. Then this normalized intensity was divided by the sum normalized intensity of the other peptides not carrying any PTMs for the same protein. The latter normalization would cancel the effect of abundance changes in the protein level upon treatment with the drug. Finally, we provided the trend of abundance changes for each modified peptides across different concentrations vs. respective controls and the results for all the PTM-carrying peptides for both dasatinib and staurosporine are shown in Supplementary Figure 5 in Supplementary data 5. While some modified peptides show a trend of decreasing or increasing abundance in a concentration-dependent manner, some other PTM-carrying peptides are unchanged upon different treatments, as expected. For example, the levels of two phosphopeptides (with three phosphothreonine) belonging to myristoylated alanine-rich C-kinase substrate (MARCKS) decreased by 1.5-fold upon treatment with the highest concentration of dasatinib. The PTM-carrying peptides following a concentration-dependent trend could be involved in drug mechanism of action.

## Discussion

In the present work, we introduce an informatic pipeline called PTM-centric proteomics for prediction of relevant PTMs using PEIMAN2 R package. This informatic pipeline does not perform *de novo* PTM identification; instead, it relies on PTM annotations from curated databases, such as UniProtKB/Swiss-Prot, to predict modifications based on existing knowledge. In developing PEIMAN2, we based our approach on the PTM keyword list from UniProtKB/Swiss-Prot. For example, during development, we removed “Nucleotide binding” from the PTM list in line with UniProtKB’s updates. For certain PTMs with ongoing debates— such as acetylation, which some researchers classify as co-translational rather than post- translational^24^—we have chosen to align with UniProtKB’s annotations to ensure consistency in our analyses. Additionally, we recognize that some modifications, such as disulfide bridges (DBBs), appear frequently in our results largely due to their overrepresentation in the PTM database. We’ll clarify that while DBBs are frequently detected due to database representation, their biological relevance should be evaluated in the context of each study’s specific focus.

To demonstrate the efficacy of this tool, we applied it to predict pertinent PTMs for differentially expressed proteins in response to perturbations or interventions. Specifically, we present a case study involving the usage of our in-house multikinase inhibitors. We also demonstrate how this tool can direct future research towards investigating a particular biologically functional PTM within a study, as evidenced by our glycoproteomics case study. Our demonstrations highlight that the inclusion of these predicted modifications in a refined database search leads to the identification of a greater number of proteins and the detection of PTMs on the most highly expressed proteins. When the answer to the research question lies beyond expression proteomics (usually does), in order to explain a particular phenotype, these findings are highly relevant and informative.

Any alterations in PTM processing in response to perturbations can potentially impact various biochemical and biophysical aspects of proteins, subsequently affecting the cellular and even organismal phenotype. For example, a simple hypusination event on eIF5A drives protein synthesis and cell proliferation^25^. Therefore, PTM studies are one of the major forefronts or proteomics research and their widespread use is not limited only to mammals or eukaryota. PTM types and sites in bacteria, archaea, and even viral proteins have been characterized and reported in an extensive number of studies^26–28^. These PTMs can modulate protein turnover^29^, stability/solubility^2, 8, 30, 31^, folding and localization of proteins, the interactions between proteins^32^, genome function^33^, the trafficking of molecules^34^, cell signaling ^35^ and the activation of receptors^36^. For instance, in our first case study, we leveraged recently published data that elucidated the influence of N-glycosylation on key clinicopathological features and patient survival in the context of oral cancer^19^. In addition, since many proteins include numerous PTMs and one PTM can change the prevalence or occupancy of others, a phenomenon known as PTM crosstalk, it is difficult to understand these complex control mechanisms without characterization and analysis of PTMs^37^.

There have been many efforts to focus on investigating the identity and effect of a single PTM or multiple PTMs on protein function^38–40^. Although mass spectrometry-based proteomics is the golden standard for PTM analyses, the high-throughput experimental procedures used to identify PTMs are labor intensive and time-consuming^41^. Suppose one has a presumption about the role of protein phosphorylation. In that case, we need to design a phosphoproteomics study using titanium/zirconium dioxide-based beads to enrich phosphopeptides with high specificity. Otherwise, the study design is independent of adding extra experimental steps and routine proteomics database searches are applied. In spite of these advances, PTM identification is not the focal point of any proteomics investigation that lacks a prior PTM-specific hypothesis. Therefore, there is an immediate demand for computational methodologies and effective tools that can predict PTMs that are most probably found in a given biological sample or are occurring upon a specific perturbation^6^.

Savitski *et al.* provided a computational method called ModifiComb based on the difference between the molecular masses and the retention time of the modified and unmodified peptides^18^. The authors provided a method that is independent of PTM-related *priori* assumptions. Compared to searching all possible modifications, this method succeeded to reduce search space and, as a result, the propensity of false positive PTM identification. Na *et al.* introduced MOD(i) as an innovative solution to address a common limitation in existing tools, namely the oversight of rare PTMs that could offer crucial insights into protein functionality^42^. Unlike ModifiComb, MOD(i) leverages tag chains to adeptly pinpoint modified regions within spectra, showcasing its effectiveness not only in identifying various PTMs within proteins but also in discerning multiply modified peptides. This approach not only enhances the comprehensive analysis of PTMs in proteins but also holds promise for uncovering novel and less common modifications that may play pivotal roles in understanding protein functions. To improve the understanding of this dark matter of proteomics, Kong *et al.* also presented a fragment-ion indexing method and implemented it into MSFragger tool to computationally speed up searching proteomics database with PTMs^43, 44^. While these methods employ a chemometric viewpoint, such as identifying unmodified peptides to compute differences in mass and retention time or using tag chains to identify modified regions within a spectrum, it’s essential to recognize that not all detectable chemical modifications in mass spectra bear biological significance. From a biological perspective, PEIMAN2 incorporates enrichment analysis after differential expression analysis to avoid detecting inert, stochastically modified peptides, predicting biologically important PTMs, termed functional PTMs. Note that the statistical methods used in PEIMAN2, including SEA and PSEA, may face limitations due to the limited annotations available in the UniProtKB/Swiss- Prot. These limitations can affect the detection of low-abundance PTMs, as certain biologically relevant modifications may remain undetected due to their subtle or rare occurrence.

However, PEIMAN2 holds considerable potential for clinical applications, particularly in advancing diagnostic and therapeutic strategies. It is well established that specific PTMs, such as phosphorylation, ubiquitination, and glycosylation, play crucial roles in disease processes, making them valuable as biomarkers for early detection or as therapeutic targets ^45–47^. Identifying and focusing on these PTMs could greatly enhance disease diagnosis and treatment approaches. For example, phosphorylation, glycosylation, and ubiquitination patterns are strongly associated with cancer, with some, like FDA-approved PTM biomarkers for glioblastoma, already applied in clinical diagnostics^48^. Aberrant glycosylation, commonly observed in oncogenic transformations, serves as an early cancer biomarker, with markers like CA19-9, CA125, and PSA used to detect and monitor cancer progression in clinical practice^49^. On the therapeutic front, PTM-targeting drugs show promise in modulating disease-critical pathways. For instance, kinase inhibitors targeting phosphorylation, such as imatinib, have significantly advanced cancer treatment, with inhibitors of MEK, PI3K, and ERK pathways also under clinical evaluation^50, 51^. Histone deacetylase inhibitors (HDACi) address tumorigenic deacetylation imbalances, while PARP inhibitors targeting PARylation pathways critical to DNA repair, offer further examples of PTM-based interventions^52^. These cases highlight the potential of PTM-centric proteomics to map disease-specific PTMs, enabling precise diagnostics and informing targeted therapeutic strategies to advance personalized medicine.

We believe that PTM-centric proteomics based on enrichment analysis is a successful attempt to bring the results of perturbational studies closer to reality. Such predictions present opportunities for developing myriad PTM-related hypotheses and a particular follow-up experimental design in biological studies. To carry out PTM-centric proteomics, it is recommended to incorporate PEIMAN2 after the initial round of mass spectrometry database search and analysis in order to carry out a second round of refined mass spectrometry database search and downstream analysis with a given set of PTMs. PEIMAN2 is not dependent on data from any MS instrument and can be easily integrated into the majority of existing data analysis pipelines. Accordingly, PEIMAN2 has the potential to become a valuable option for routine analysis of the most probable PTMs in shotgun proteomics data. Furthermore, the application of this software package is expected to enhance the exploration of PTM crosstalk in the contexts of both homeostasis and disease.

## Methods

### Cell culture

Human lung carcinoma A549 were grown in McCoy’s 5A medium, supplemented with 10% FBS superior (Biochrom, Berlin, Germany), 2 mM L-glutamine (Lonza, Wakersville, MD, USA) and 100 units/mL penicillin/streptomycin (Gibco, Invitrogen) and incubated at 37 °C in 5% CO2. Cells were routinely checked for mycoplasma contamination by PCR and low passage number cells from ATCC were used in the experiments.

### Cell viability assay

Cell viability upon compound treatment was measured using CellTiter-Blue assay (Promega) according to manufacturer protocol and the LC50s were determined as the concentration of compound causing 50% cytotoxicity.

### Multikinase inhibitor treatment

After seeding 250,000 A549 cells in triplicates in 6 well plates, cells were allowed to grow for 24 h, after which they were treated with the compounds for 24 h in triplicates. Dasatinib was profiled at 100 nM, 1 *µ*M, 5 *µ*M and 25 *µ*M. Staurosporine was profiled at 8 nM, 40 nM, 200 nM and 1 *µ*M. Cells treated with DMSO were used as controls.

**Table 1:**
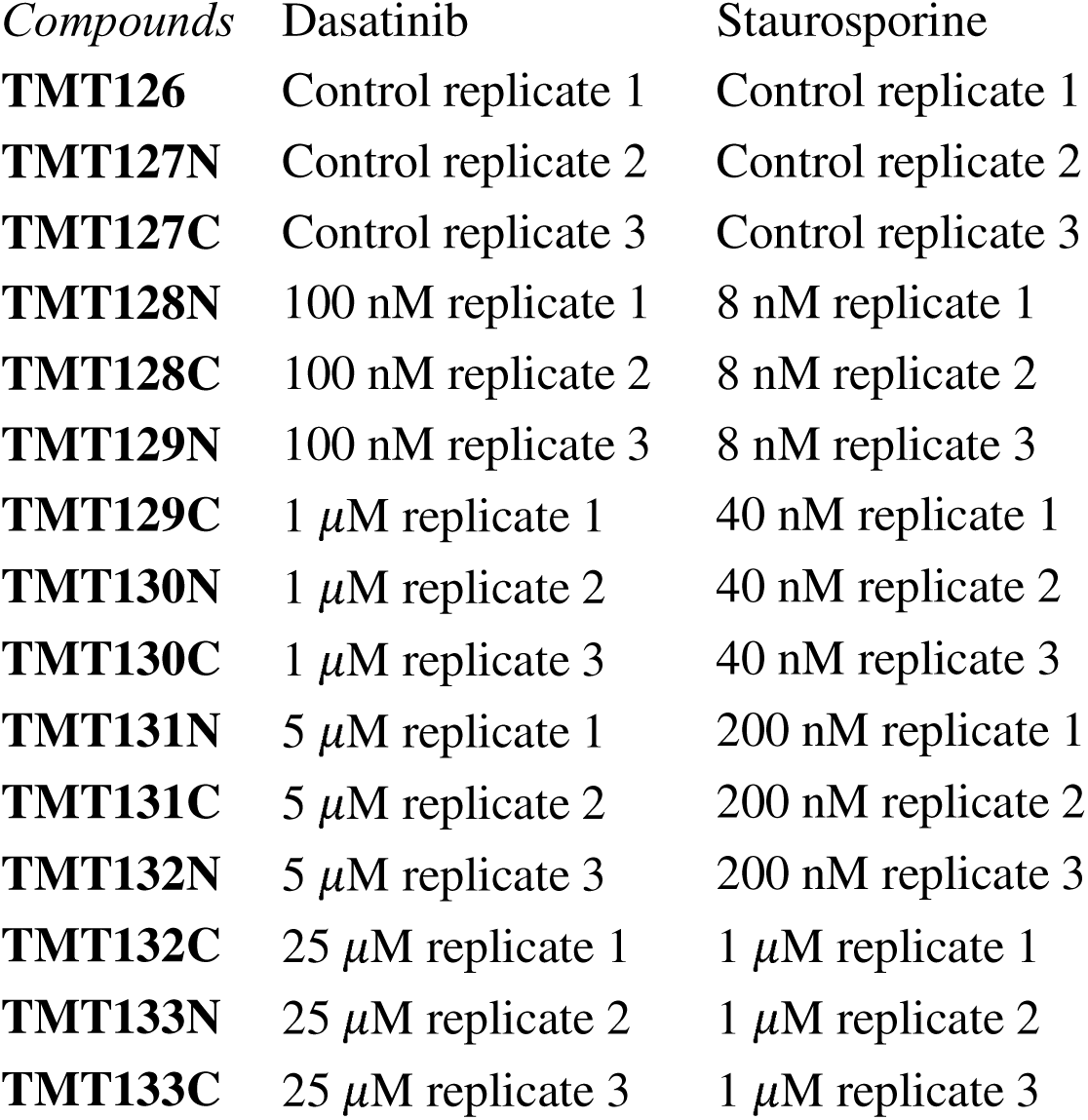
TMT labeling scheme for the experiments.

| Compounds | Dasatinib | Staurosporine |
| --- | --- | --- |
| **TMT126** | Control replicate 1 | Control replicate 1 |
| **TMT127N** | Control replicate 2 | Control replicate 2 |
| **TMT127C** | Control replicate 3 | Control replicate 3 |
| **TMT128N** | 100 nM replicate 1 | 8 nM replicate 1 |
| **TMT128C** | 100 nM replicate 2 | 8 nM replicate 2 |
| **TMT129N** | 100 nM replicate 3 | 8 nM replicate 3 |
| **TMT129C** | 1 μM replicate 1 | 40 nM replicate 1 |
| **TMT130N** | 1 μM replicate 2 | 40 nM replicate 2 |
| **TMT130C** | 1 μM replicate 3 | 40 nM replicate 3 |
| **TMT131N** | 5 μM replicate 1 | 200 nM replicate 1 |
| **TMT131C** | 5 μM replicate 2 | 200 nM replicate 2 |
| **TMT132N** | 5 μM replicate 3 | 200 nM replicate 3 |
| **TMT132C** | 25 μM replicate 1 | 1 μM replicate 1 |
| **TMT133N** | 25 μM replicate 2 | 1 μM replicate 2 |
| **TMT133C** | 25 μM replicate 3 | 1 μM replicate 3 |

### LC-MS/MS sample preparation

Sample preparation was done according to our previous protocol^53^. After treatment, cells were trypsinized, washed with PBS and lysed with the lysis buffer (8 M urea, 1% SDS, 50 mM Tris pH 8.5). Protein concentration was measured using Pierce BCA Protein Assay Kit (Thermo), and the volumes corresponding to 25 µg of protein was transferred from each sample to new low-bind Eppendorf tubes. DTT was added to a final concentration of 10 mM and samples were incubated for 1 h at room temperature. Subsequently, iodoacetamide (IAA) was added to a final concentration of 50 mM and samples were incubated at room temperature for 1 h in the dark. The reaction was quenched by adding an additional 10 mM of DTT. After precipitation of proteins using methanol/chloroform, the semi-dry protein pellets were dissolved in 25 µL of 8 M urea in 20 mM EPPS (pH 8.5) and were then diluted with EPPS buffer to reduce urea concentration to 4 M. Lysyl Endopeptidase (Wako) was added at a 1:75 w/w ratio to protein and incubated at room temperature overnight. After diluting urea to 1 M, trypsin (Promega) was added at the ratio of 1:75 w/w and the samples were incubated for 6 h at room temperature. TMT reagents were added 4x by weight to each sample, followed by incubation for 2 h at room temperature. The reaction was quenched by addition of 0.5% hydroxylamine. Samples were combined, acidified by TFA, cleaned using Sep-Pak (Waters) and dried using a DNA 120 SpeedVac concentrator (Thermo). Samples were resuspended in 20 mM ammonium hydroxide and separated into 96 fractions on an XBrigde BEH C18 2.1x150 mm column (Waters; Cat#186003023), using a Dionex Ultimate 3000 2DLC system (Thermo Scientific) over a 48 min gradient of 1-63% B (B=20 mM ammonium hydroxide in acetonitrile) in three steps (1-23.5% B in 42 min, 23.5-54% B in 4 min and then 54-63%B in 2 min) at 200 *µ*L/min flow. Fractions were then concatenated into 24 samples in sequential order (e.g. A1, C1, E1 and G1).

### Proteomics

After resuspension in 0.1% FA (Fluka), fractions (1 *µ*g) were analyzed by LC-MS/MS. Samples were loaded onto a 50 cm column (EASY-Spray, 75 *µ*m internal diameter (ID), PepMap C18, 2 *µ*m beads, 100 Å pore size) connected to a nanoflow Dionex UltiMate 3000 UHPLC system (Thermo) and eluted in an organic solvent gradient increasing from 4% to 26% (B: 98% ACN, 0.1% FA, 2% H2O) at a flow rate of 300 nL/min over a total 110 min method time. The eluent was ionized by electrospray and mass spectra of the molecular ions were acquired with an Orbitrap Fusion mass spectrometer (Thermo Fisher Scientific) in data-dependent mode at MS1 resolution of 120,000 and MS2 resolution of 60,000, in the m/z range from 400 to 1600. Peptide fragmentation was performed via higher-energy collision dissociation (HCD) with energy set at 35 NCE and MS2 isolation width at 1.6 Th.

### Proteomic Data, Bioinformatic and statistical Analysis

The raw LC-MS data were analyzed MaxQuant version 2.5.0.0 ^54^. The Andromeda search engine ^55^ was run against the UniProtKB/Swiss-Prot database (human version 9606, release Dec 2023). Methionine oxidation was selected as variable modifications, while cysteine carbamidomethylation was set as a fixed modification. No more than two missed cleavages were allowed, and a 1% FDR was used as a filter at both protein and peptide levels. The precursor ion and fragment ion mass tolerances in the database search were set to 4.5 ppm and 20 ppm, respectively. All the contaminants were removed in the first step. After PEIMAN2 analysis, the mentioned modifications were added to the database search. The maximum number of modifications per peptide was set to 5. All the experiments were performed in triplicates.

### Glycoproteomics data source

The dataset utilized in this study was derived from Carnielli *et al.*^19^ and is publicly accessible. It encompasses (glyco)peptide profiling data acquired through liquid chromatography-tandem mass spectrometry (LC-MS/MS). This dataset includes information from 31 patients diagnosed with OSCC, categorized into two groups with and without lymph node metastasis. The glycoproteomics and proteomics datasets underwent comprehensive analyses through Byonic and MaxQuant software, respectively. These analyses adhered to rigorous statistical methodologies and quality control protocols, yielding a dataset that offers valuable insights into the glycoproteomic and proteomic profiles of OSCC patients. Subsequently, we leveraged differentially expressed proteins and glycoproteins as input for conducting enrichment analysis within the PEIMAN2 software.

### Experimental Design and Statistical Rationale

To assess multikinase inhibitors via mass spectrometry (MS), three control samples (A549 cells treated with DMSO), three samples treated with dasatinib, and three samples treated with staurosporine were submitted to the respective platform. For drug treatments, we employed all biological replicates of independent WT cultures (n = 3) across four different concentrations, as outlined in the preceding sections. Additionally, three technical replicates accompanied each biological replicate for mass spectrometry runs.

MS data underwent validation with a 1% false discovery rate threshold for peptide identifications. Subsequent assessment of statistical significance involved a Student’s t-test. Utilizing this method, we identified 863 proteins with significant adjusted p-values less than 0.05, including 33 proteins with an absolute value of Log2FC greater than 0.6 for Dasatinib. The numbers are 817 proteins with significant adjusted p-values less than 0.05 and 57 proteins with an absolute value of Log2FC greater than 0.6 for Staurosporine. The MS proteomics data have been deposited in the ProteomeXchange Consortium via the jPOST partner repository, with the dataset identifiers PXD037679 and PXD037681.

For Glycoproteomics analysis, we leveraged publicly available data from Carnielli et al. 19 Their study included 31 OSCC patients, of whom 19 had lymph node metastasis (N+), and 12 did not (N0). Statistical significance in their analysis was determined using unpaired two-tailed Student’s t-tests, with a confidence threshold of p < 0.05.

### PEIMAN2 R package

At the first step, we downloaded 569,794 “Reviewed (UniProtKB/Swiss-ProtKB/Swiss- Prot), Manually annotated” proteins (as of September 2023 from UniProtKB/Swiss-Prot online repository, available at https://www.uniprot.org). The database records various useful functional information about proteins. We reduced the size of file by narrowing each protein record to include unique UniProtKB accession code (AC), organism taxonomy name (OS), keywords (KW) and features (FT). We were particularly interested in KW and FT as any manually curated information regarding PTM are available in these fields. At the next step, we used R software to build a database to obtain PTM profile of proteins for all species. In our search for PTMs in proteins, we used a list of controlled PTM vocabulary provided by UniProtKB/Swiss-Prot (available at https://ftp.uniprot.org/pub/databases/uniprot/current_release/knowledgebase/complete/docs/ptmli st.txt). To find proteins with any PTM modification, we searched the downloaded file against controlled PTM vocabulary by looking into KW and FT entry of each protein. The presence of any PTM in CROSSLNK, LIPID, or MOD_RES feature of any protein was of interest. At the time of preparing this manuscript, we obtained the PTM profile of 134,783 proteins. To keep up with monthly changes in UniProtKB/Swiss-Prot, we automated the preparation process and will update the database each month accordingly. The latest PEIMAN2 scripts and PEIMAN2 database are available at JafariLab GitHub repository (https://github.com/jafarilab/PEIMAN2). In addition to its functionality in the R environment, a PEIMAN2 plugin has been developed for the Perseus software platform, further enhancing accessibility for users. This plugin is available in the JafariLab GitHub repository: https://github.com/jafarilab/PEIMAN2plugin and the Perseus Plugin Store: https://www.maxquant.org/perseus_plugins/.

### Enrichment Analysis

#### Singular Enrichment Analysis (SEA)

The enrichment analysis is a powerful strategy which facilitates the identification of biological processes for a list of genes or proteins. The SEA is known as one of the traditional methods to infer the biological functions in a given list of genes. The analysis in SEA starts with a list of differentially expressed genes provided by researcher (selected with some criteria: p-value or fold-change). The idea behind SEA is to test if the number of genes in the list with a certain biological function (for example PTM) is significantly different from occurrence through random chance. In a general sense, enrichment analysis investigates whether a group of genes or proteins are over/under-presented for a specific biological pathway in a large set of genes/proteins. Different statistical methods are introduced to measure this discrepancy such as Chi-Square, Fisher’s exact test, and hypergeometric test. We previously implemented a standalone software to run SEA in a list of proteins and infer any enriched modification by applying a hypergeometric test^18^. The idea in PEIMAN standalone software and PEIMAN2 is to investigate if a subset of proteins is over/under presented for any particular PTM, in a large set of proteins. We here briefly describe the idea of hypergeometric test in this context. Assume there are *N* proteins in the database and *K(≤N)* of these proteins have one of the known modifications, for example “Acetylation”. We pass a list of *n* proteins. We can apply hypergeometric test to check if “Acetylation” is over-under/represented in the sample list of *n* proteins using a hypergeometric test. The p-value of such a test is calculated as:

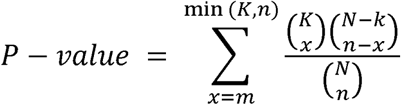

This simple idea is very helpful in inferring biological meaning from a large list of genes. In the next section, we discuss some of the weaknesses associated with SEA and review an alternative powerful approach to SEA.

### Protein Set Enrichment Analysis (PSEA)

SEA takes a list of differentially expressed genes/proteins and identifies if there is a significant overrepresentation of genes or proteins associated with a specific biological feature. However, SEA has known limitations that warrant consideration^56^. Firstly, when correcting for multiple testing, the resulting p-values may render no genes or proteins differentially significant, particularly when real differences are subtle compared to inherent noise. Secondly, the final list generated by SEA may contain a multitude of declared significant genes or proteins, which can pose challenges in interpretation and may lead to subjective interpretations among biologists with varying expertise. Thirdly, single gene or protein enrichment analysis has the potential to overlook important effects on pathways. Lastly, it is not uncommon for different research laboratory groups to report multiple lists of significant genes or proteins for the same perturbation or biological process.

Gene set enrichment analysis or GSEA has been introduced by Subramanian *et al*. ^56^ to overcome these drawbacks. We briefly highlight the key points of GSEA method here. GSEA is applied on profiles of genome-wide expression data that belongs to two experimental groups (control vs treatment). Genes are then sorted based on a given score, for example correlation between their expression profile or the class they belong to. A set of genes, S, is defined as genes that belong to a certain set with a distinct biological annotation (e.g., metabolic pathway, GO category, PTM). The idea behind GSEA is to identify if the members of set S tend to show more often toward the top (or bottom) of the gene list or are randomly distributed throughout the list.

The GSEA method can be summarized in three steps as follows. First, an enrichment score (ES) is calculated to measure if the gene set S is over-presented at the top or bottom of the ranked gene list. This is achieved by calculating a signed version of the Kolmogorov-Smirnov statistic while running from the top to the bottom of the ranked list. Whenever we encounter a gene in the S set, the value of the statistic is increased proportional to an exponent power of the gene’s score. Likewise, when the gene is not in the S set, the value of statistic is decreased. The enrichment score for gene set S is defined as the maximum observed deviation from zero in the running score profile. In the second step, the significance of calculated ES score for a given list of genes is evaluated by randomly permuting the score of each gene for certain number of times (usually 1000 times) and calculating ES for each random profile to generate a null distribution for the ES. A p-value based on the permutations is calculated according to the null distribution. Finally, GSEA accounts for the effect of multiple testing by calculating FDR. This is achieved by normalizing ES score of each gene set relative to gene set size. For more details refer to supplementary material of ^31^.

Inspired by the idea and usefulness of GSEA in elucidating biological inferences in a given list of genes, we implement protein set enrichment analysis or PSEA in PEIMAN2 package to infer biological meaning from a list of proteins. In our work, the gene sets are replaced by a set of proteins that belong to a certain modification group, for example “Acetylation”. For any list of protein given by researcher, the set of proteins with a certain modification are identified. For each set of proteins, we calculate enrichment score and assess the significance of ES by the methods described in ^43^. Finally, we provide a list of modification that are most probably enriched in a given list. All these functionalities are implemented in an R package to serve a broader community of researchers. For more details on functionality of package, please read the Vignette and Readme page at PEIMAN2 GitHub page.

## Data availability

The LC-MS/MS raw data files and extracted peptides and protein abundances are deposited in the jPOST repository of the ProteomeXchange Consortium ^57^ under the dataset identifiers PXD037679 and PXD037681. Additionally, the LC-MS/MS raw data files for the glycoproteomics case study are available through the ProteomeXchange Consortium via the PRIDE repository under the dataset identifier PXD037134.

## Code availability

All analyses reported in this study used the statistical software R (v.4.0.0). The R package can be found on GitHub (https://github.com/jafarilab/PEIMAN2).

## Supporting information

Supp5Combin

## Acknowledgements

This study was financially supported by the Research Council of Finland [Grant 332454 to M.J.], Jane and Aatos Erkko foundation [Grant 220031 to M.J.], Swedish Research Council [Grant 2020-00687 to A.A.S.], and the Swedish Society of Medicine [Grant SLS-961262, 1086 Stiftelsen Albert Nilssons forskningsfond to A.A.S.]. We would like to thank Prof. Roman A. Zubarev for his valuable comments and input in the manuscript. Meilahti Clinical Proteomics Core Unit is supported by Biocenter Finland and Helsinki Institute of Life Sciences (HiLIFE).

## Author information

### Contributions

M.J. conceived of the study and supervised the project. P.N. and M.J. developed the computational analysis as well as the PEIMAN2 R package. U.V. developed PEIMAN2 Perseus plugin. A.A.S designed, developed, and led the experimental methods for deep expression profiling. M.M., A.A.S, and M.J. contributed to the interpretation of the findings, and M.B. advised on the work. All authors contributed to the final manuscript by discussing the findings and reviewing and modifying it.

## Supplementary data

**Supplementary data 1**. A table of all identified proteins for dasatinib following PTM-centric proteome informatic pipeline. This file represents the raw output from MaxQuant search before undergoing any filtering for subsequent analysis.

**Supplementary data 2.** A table of all identified proteins for staurosporine following PTM-centric proteome informatic pipeline. This file represents the raw output from MaxQuant search before undergoing any filtering for subsequent analysis.

**Supplementary data 3.** A compiled table of all PTM-carrying peptides for dasatinib and calculated final fold changes that are plotted in Supp figure 5.

**Supplementary data 4.** A compiled table of all PTM-carrying peptides for staurosporine and calculated final fold changes that are plotted in Supp figure 5.

**Supplementary data 5:** This pdf file contains the Supplementary Figures 1-5.

**Supplementary Figure 1:** PTM frequency treemaps for eight popular organisms from diverse taxonomic branches of life.

**Supplementary Figure 2:** The t-SNE plots based on PTM profiles in four super kingdoms of life, i.e., Archaea, Bacteria, Eukaryota and Viruses (panels A-D). Each dot in the plots presents one organism. The red and grey colors indicates if the point (organism) belongs to corresponding super kingdom of life or not. Note that we anticipate a more uniform distribution of viruses across all three of the other phyla of life (Panel D).

**Supplementary Figure 3:** The running score plot of the first top five PTMs identified on differentially expressed proteins upon dasatinib treatment. The x-axis is the ranked protein based on their score and the y-axis is their enrichment score. The rug in the x-axis indicates the proteins with the corresponding PTM. The position of maximum running enrichment score is denoted by a red dashed line.

**Supplementary Figure 4:** The running score plot of the first top five PTMS identified on differentially expressed proteins upon staurosporine treatment. The x-axis is the ranked protein based on their score and the y-axis is their enrichment score. The rug in the x-axis indicates the proteins with the corresponding PTM. The position of maximum running enrichment score is denoted by a red dashed line.

**Supplementary Figure 5:** The modified peptides with probabilities. The modified peptides with all above-mentioned PTMs are listed separately upon dasatinib and staurosporine treatment. The x-axis is the four drug concentrations and the control, and the y-axis is the proportional abundance of the corresponding peptide compared to unmodified peptide. Note that identical quantification values may appear for peptides represented with different probabilities for different phosphorylation site localizations (see Supplementary data 3 and 4 for more details).

