## Supplementary material for "Monitoring Functional Post-Translational Modifications Using a Data-Driven Proteome Informatic Pipeline": Supp5Combin

#### Supplementary data 5

### Monitoring Functional Post-Translational Modifications Using a Data-Driven Proteome Informatic Pipeline


Payman Nickchi[1], Uladzislau Vadadokhau[2], Mehdi Mirzaie[3], Marc Baumann[2], Amir A. Saei [4,ϙ] and Mohieddin Jafari[2,ϙ,*]

[1]Department of Statistics and Actuarial Science, Simon Fraser University, British Columbia

[2]Medicum, Department of Biochemistry and Developmental Biology, Meilahti Clinical Proteomics Core Facility, University of Helsinki, Helsinki, Finland

[3]Department of Pharmacology, Faculty of Medicine & Helsinki Institute of Life Science, University of Helsinki, Helsinki, Finland

[4]Department of Microbiology, Tumor and Cell Biology, Karolinska Institutet, Stockholm, Sweden

[ϙ]These authors contributed as co-last authors.

* Corresponding author:

Mohieddin Jafari,

#### Supplementary Figures

***Supplementary Figure 1:*** *PTM frequency treemaps for eight popular organisms from diverse taxonomic branches of life.*

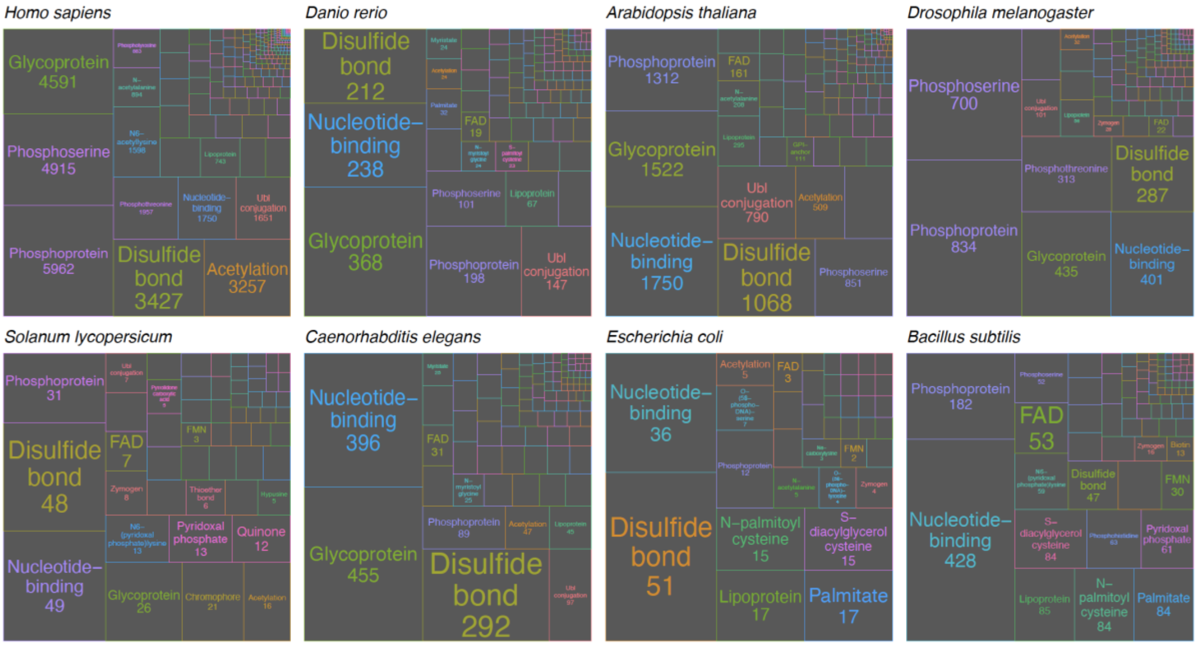


***Supplementary Figure 2:*** *The t-SNE plots based on PTM profiles in four super kingdoms of life, i.e., Archaea, Bacteria, Eukaryota and Viruses (panels A-D). Each dot in the plots presents one organism. The red and grey colors indicates if the point (organism) belongs to corresponding super kingdom of life or not. Note that we anticipate a more uniform distribution of viruses across all three of the other phyla of life (Panel D).*

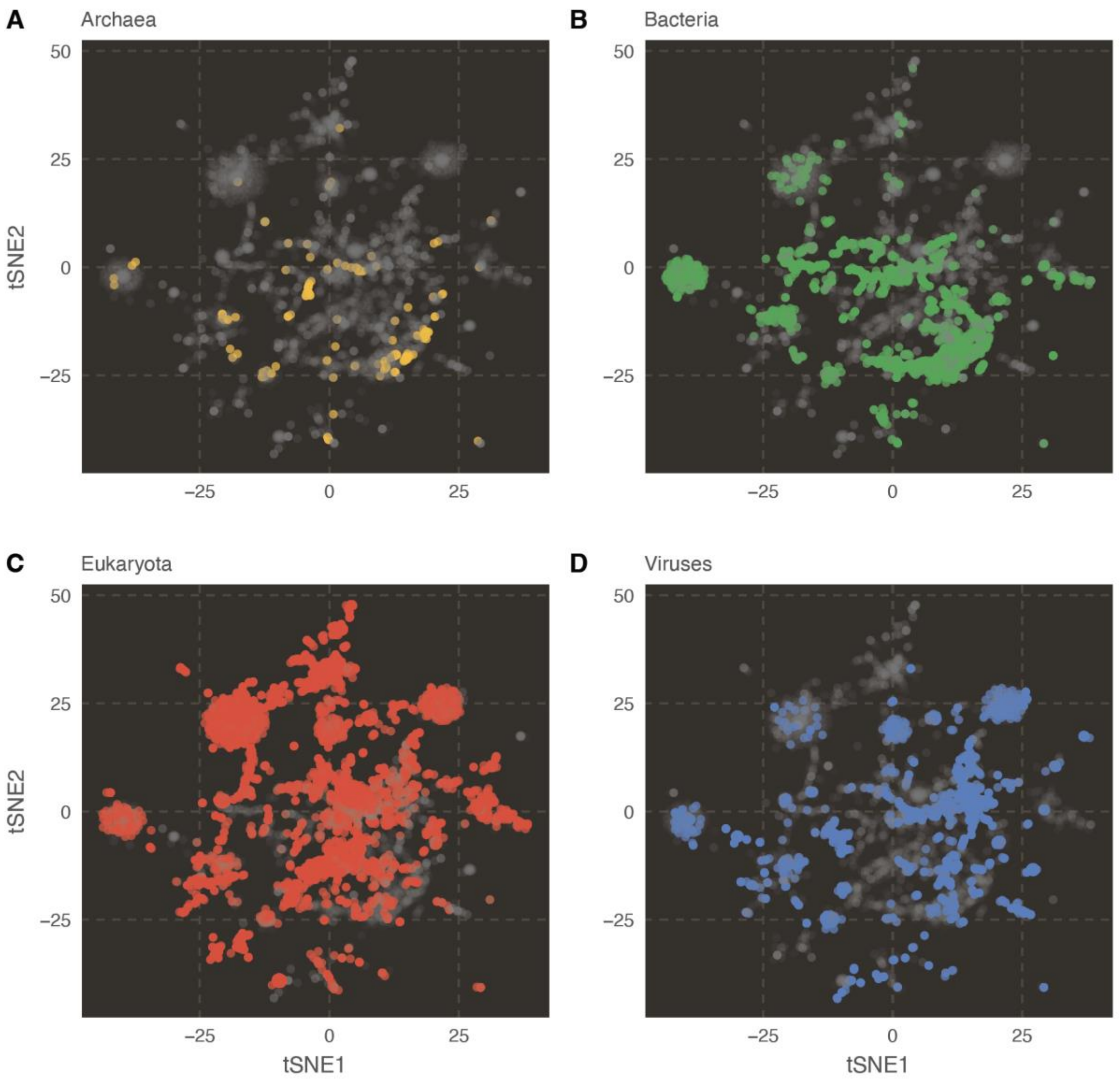

A
Archaea
B
Bacteria
C
Eukaryota
D
Viruses
tSNE2
tSNE1
50
25
0
–25

***Supplementary Figure 3:*** *The running score plot of the first top five PTMs identified on differentially expressed proteins upon dasatinib treatment. The x-axis is the ranked protein based on their score and the y-axis is their enrichment score. The rug in the x-axis indicates the proteins with the corresponding PTM. The position of maximum running enrichment score is denoted by a red dashed line.*

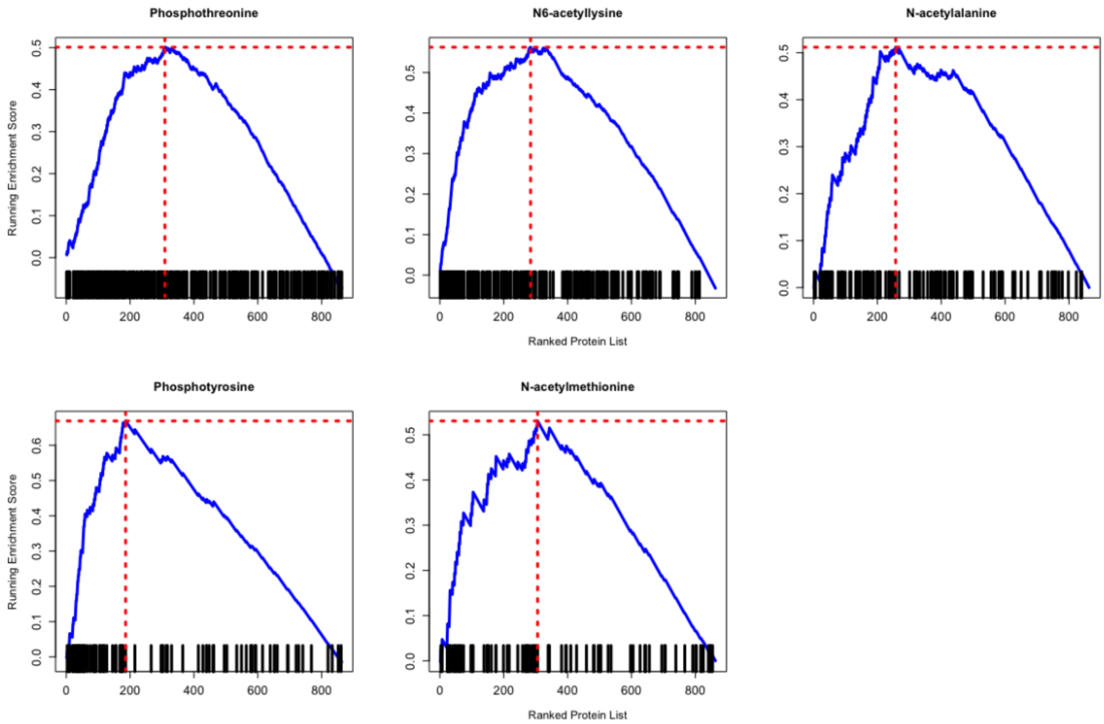

***Supplementary Figure 4:*** *The running score plot of the first top five PTMS identified on differentially expressed proteins upon staurosporine treatment. The x-axis is the ranked protein based on their score and the y-axis is their enrichment score. The rug in the x-axis indicates the proteins with the corresponding PTM. The position of maximum running enrichment score is denoted by a red dashed line.*

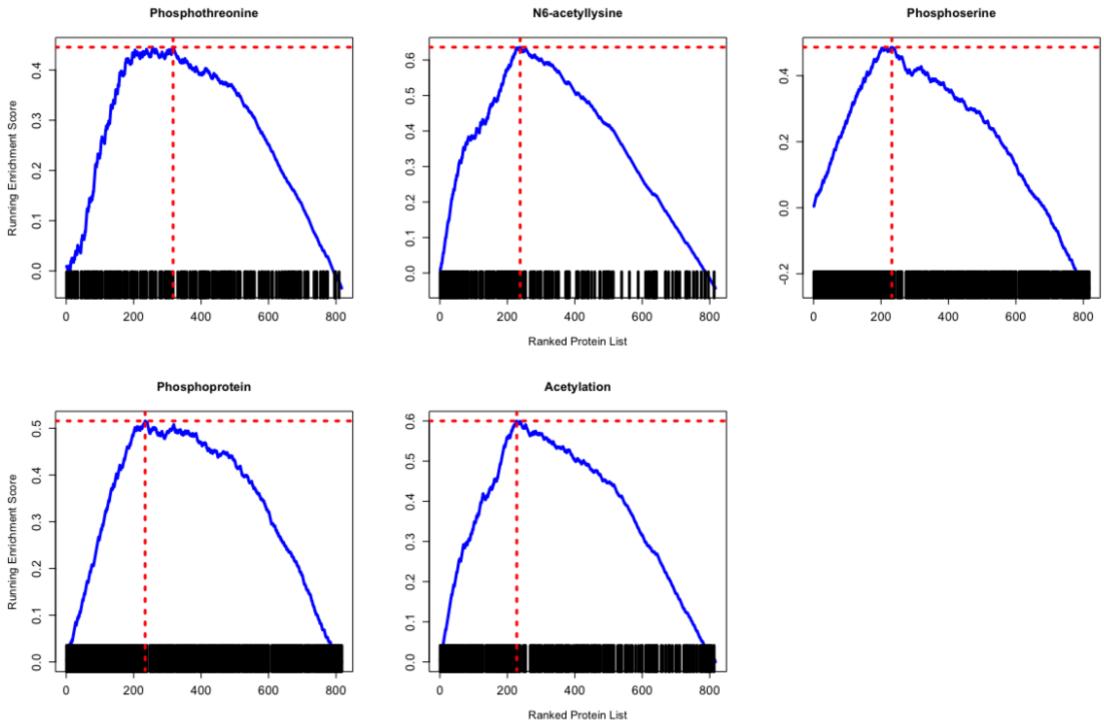

***Supplementary Figure 5:*** *The modified peptides with probabilities. The modified peptides with all above-mentioned PTMs are listed separately upon dasatinib and staurosporine treatment. The x-axis is the four drug concentrations and the control, and the y-axis is the proportional abundance of the corresponding peptide compared to unmodified peptide (see Supplementary data 3 and 4 for more details). Note: Identical peptide sequences may appear with different phosphorylation site probabilities, representing varying confidence levels in site localization.*

Dasatinib_Phospho (TY) sites with probabilities

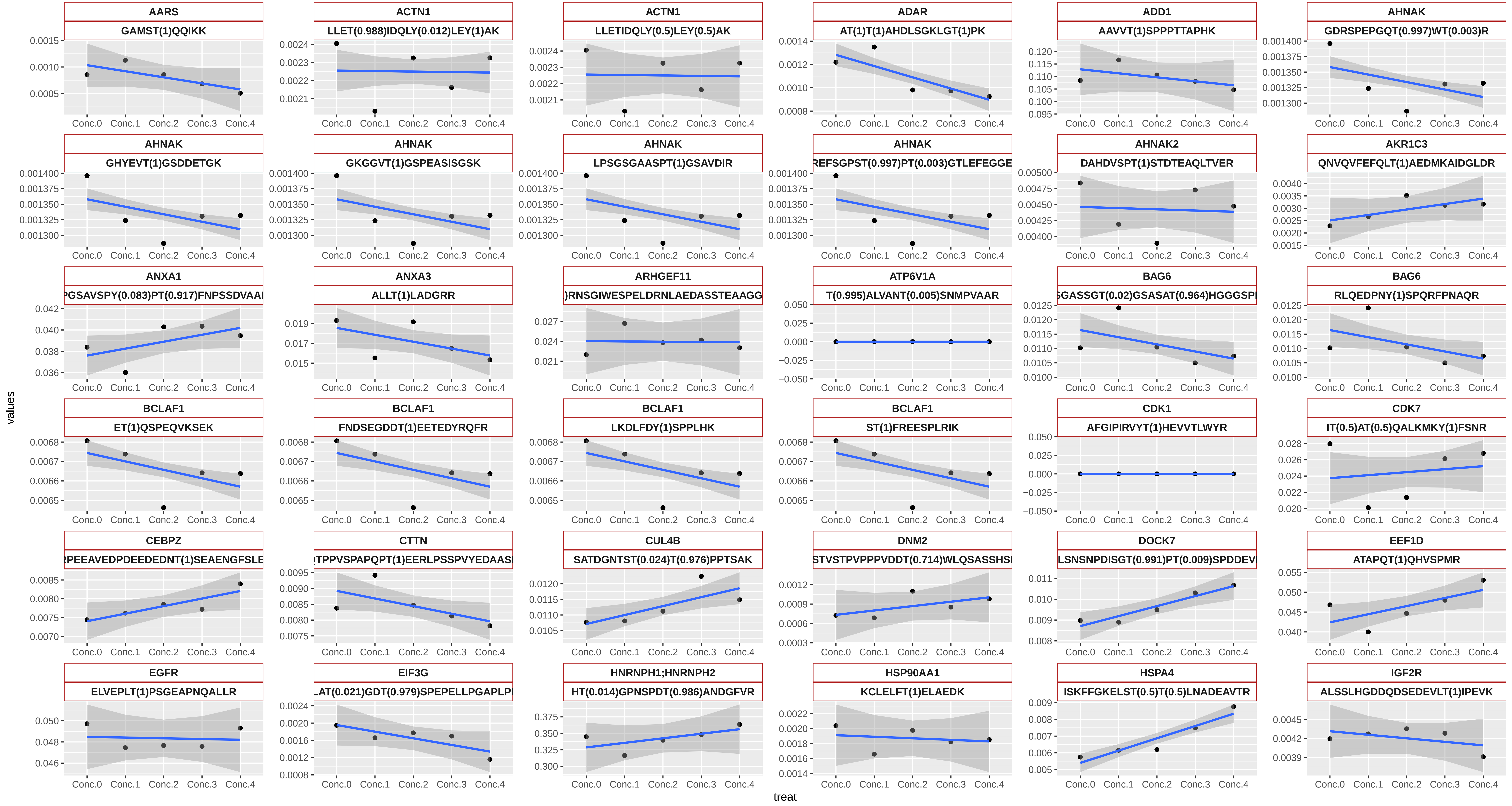

Dasatinib_Phospho (TY) sites with probabilities

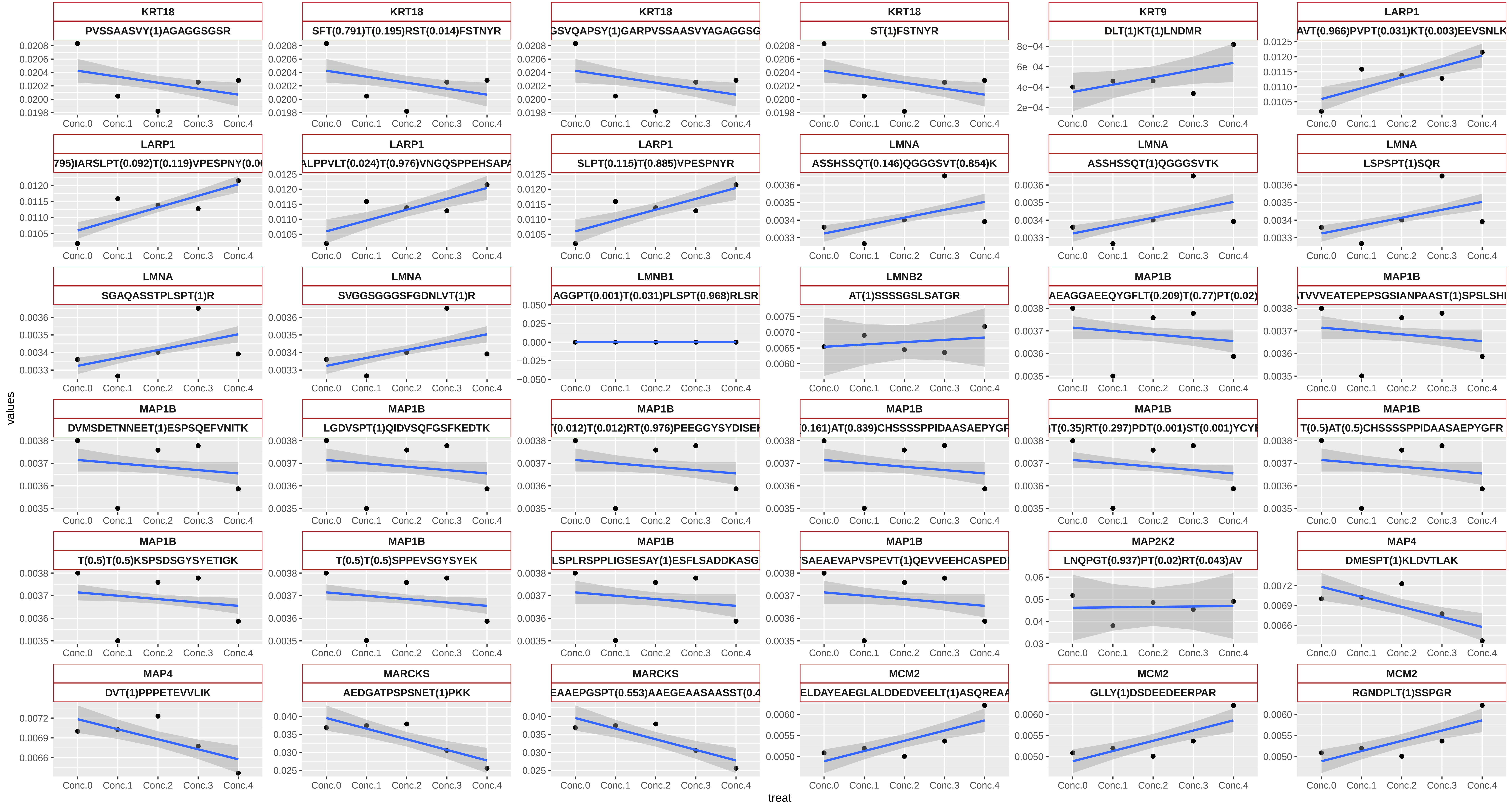

Dasatinib_Phospho (TY) sites with probabilities

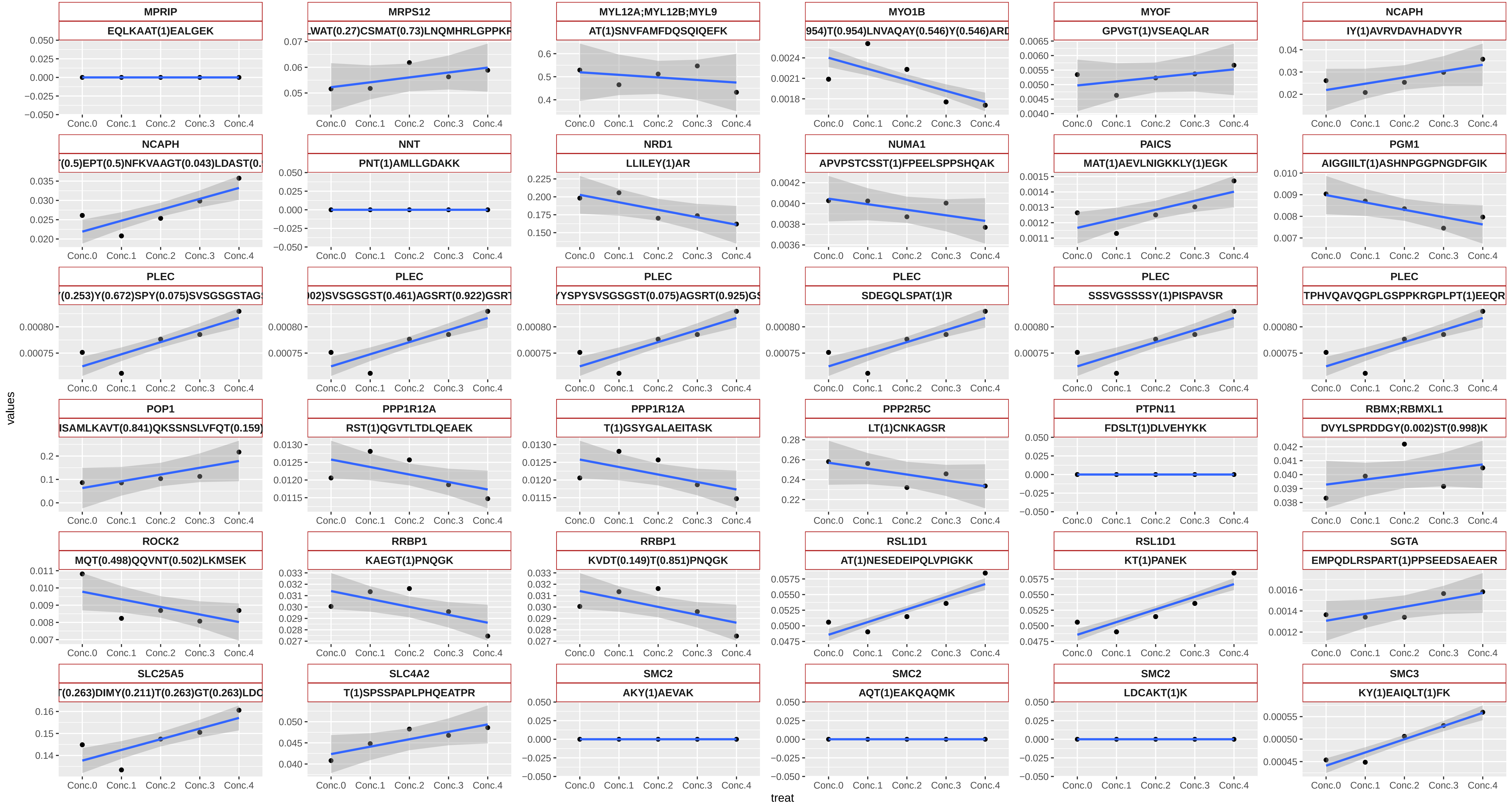

Dasatinib_Phospho (TY) sites with probabilities

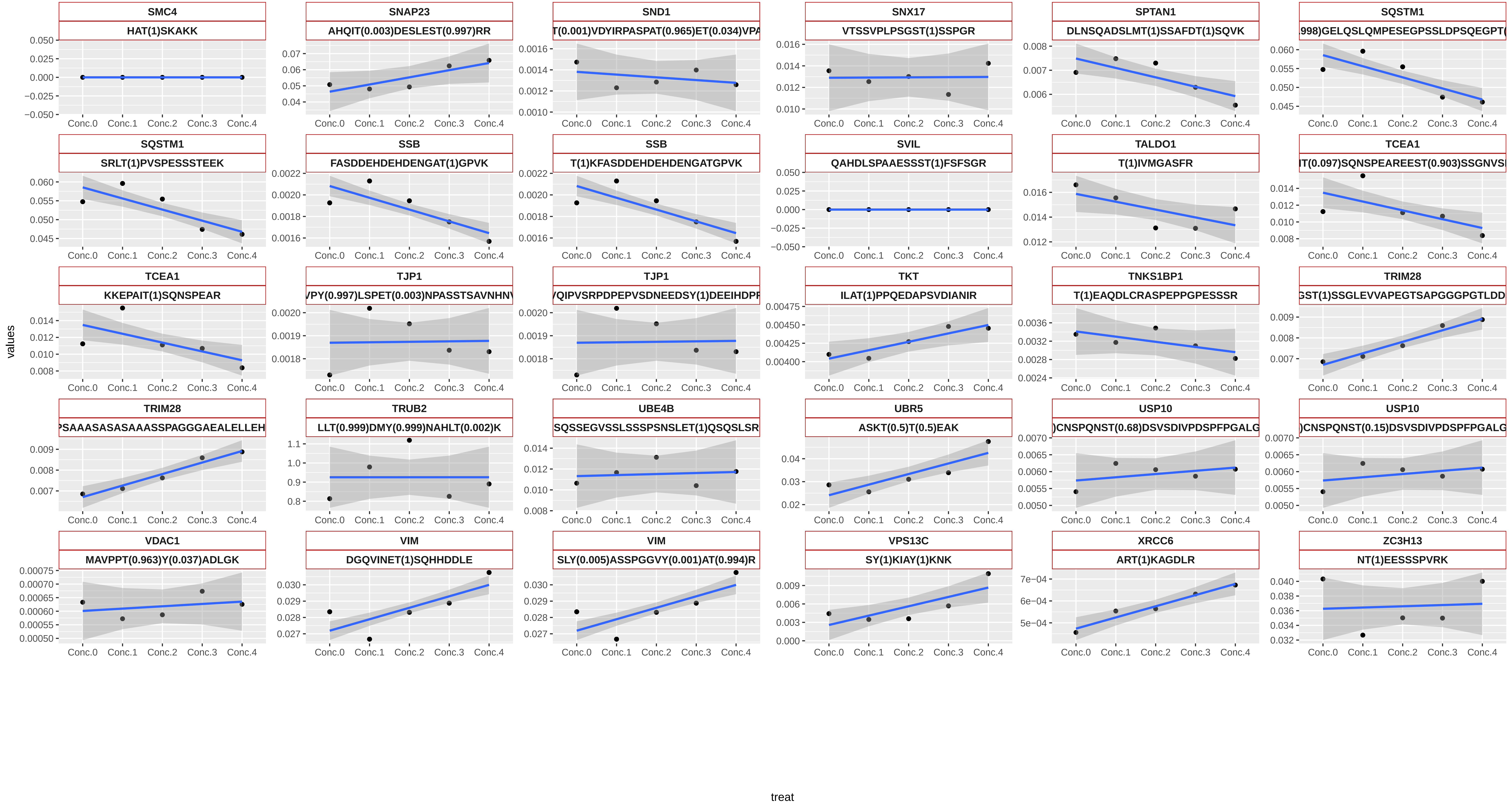

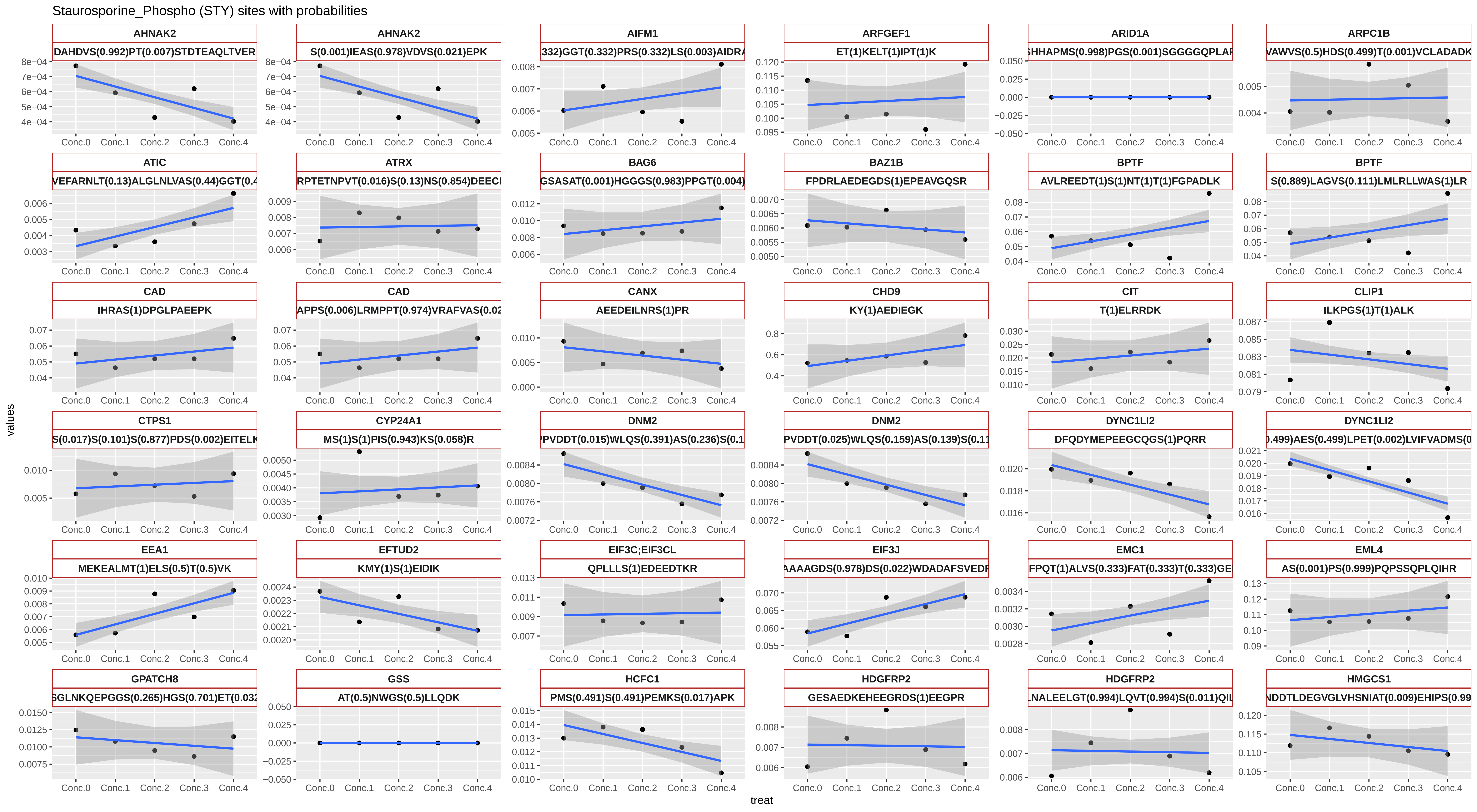
Staurosporine_Phospho (STY) sites with probabilities
AHNAK2
DAHDVS(0.992)PT(0.007)STDTEAQLTVER
AHNAK2
S(0.001)IEAS(0.978)VDVS(0.021)EPK
AIFM1
332)GGT(0.332)PRS(0.332)LS(0.003)AIDRA
ARFGEF1
ET(1)KELT(1)IPT(1)K
ARID1A
SHHAPMS(0.998)PGS(0.001)SGGGGQPLAF
ARPC1B
VAWVS(0.5)HDS(0.499)T(0.001)VCLADADK
ATIC
VEFARNLT(0.13)ALGLNLVAS(0.44)GGT(0.4
ATRX
RPTETNPVT(0.016)S(0.13)NS(0.854)DEECI
BAG6
GSASAT(0.001)HGGGS(0.983)PPGT(0.004)
BAZ1B
FPDRLAEDEGDS(1)EPEAVGQSR
BPTF
AVLREEDT(1)S(1)NT(1)T(1)FGPADLK
BPTF
S(0.889)LAGVS(0.111)LMLRLLWAS(1)LR
CAD
IHRAS(1)DPGLPAEEPK
CAD
APPS(0.006)LRMPPT(0.974)VRAFVAS(0.02
CANX
AEEDEILNRS(1)PR
CHD9
KY(1)AEDIEGK
CIT
T(1)ELRRDK
CLIP1
ILKPGS(1)T(1)ALK
CTPS1
S(0.017)S(0.101)S(0.877)PDS(0.002)EITELK
CYP24A1
MS(1)S(1)PIS(0.943)KS(0.058)R
DNM2
PPVDDT(0.015)WLQS(0.391)AS(0.236)S(0.1
DNM2
PVDDT(0.025)WLQS(0.159)AS(0.139)S(0.11
DYNC1LI2
DFQDYMEPEEGCQGS(1)PQRR
DYNC1LI2
.499)AES(0.499)LPET(0.002)LVIFVADMS(0
EEA1
MEKEALMT(1)ELS(0.5)T(0.5)VK
EFTUD2
KMY(1)S(1)EIDIK
EIF3C;EIF3CL
QPLLLS(1)EDEEDTKR
EIF3J
AAAAGDS(0.978)DS(0.022)WDADAFSVEDF
EMC1
FPQT(1)ALVS(0.333)FAT(0.333)T(0.333)GE
EML4
AS(0.001)PS(0.999)PQPSSQPLQIHR
GPATCH8
SGLNKQEPGGS(0.265)HGS(0.701)ET(0.032
GSS
AT(0.5)NWGS(0.5)LLQDK
HCFC1
PMS(0.491)S(0.491)PEMKS(0.017)APK
HDGFRP2
GESAEDKEHEEGRDS(1)EEGPR
HDGFRP2
LNALEELGT(0.994)LQVT(0.994)S(0.011)QIL
HMGCS1
NDDTLDEGVGLVHSNIAT(0.009)EHIPS(0.99
values
treat

Staurosporine_Phospho (STY) sites with probabilities

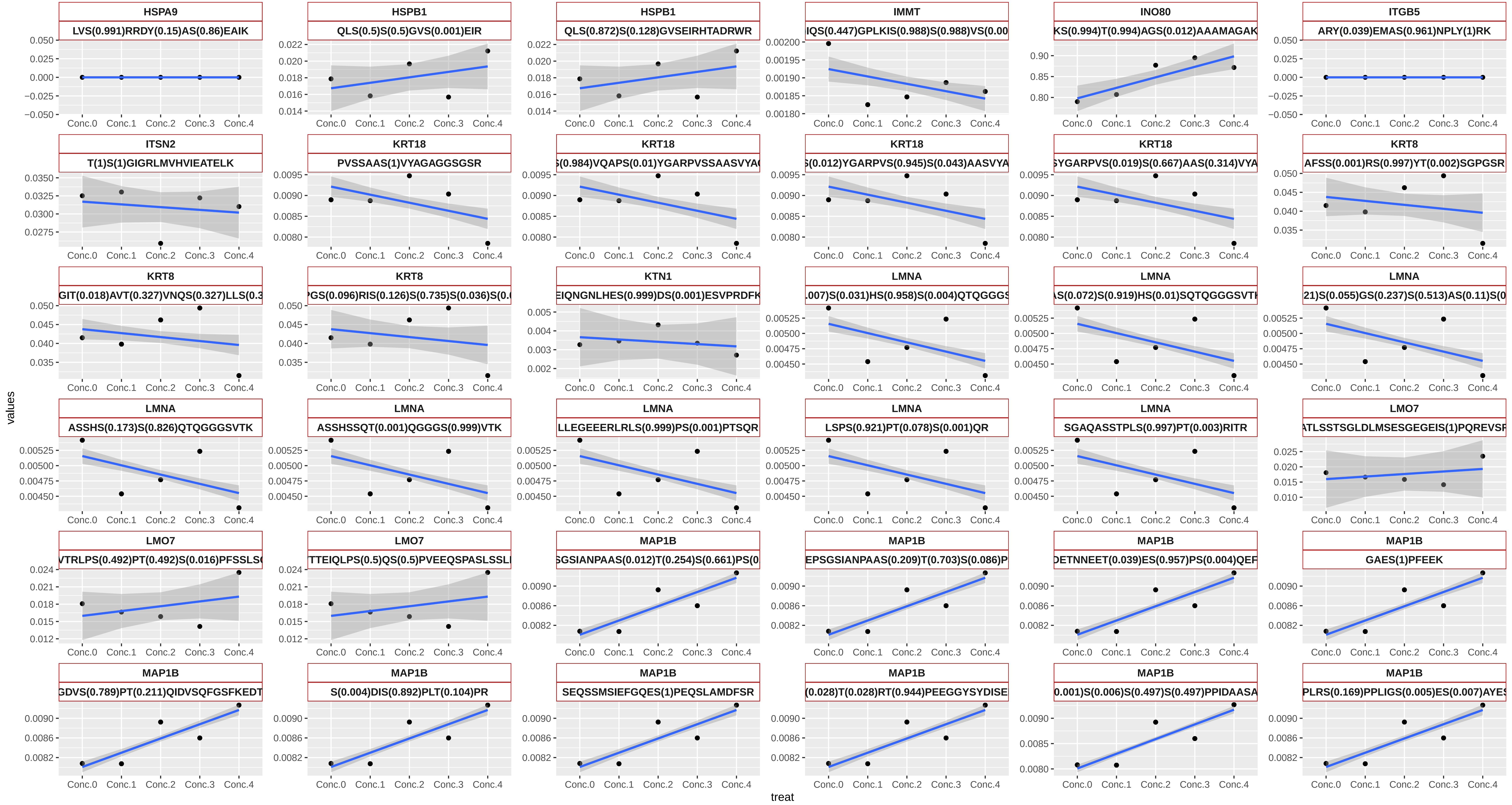

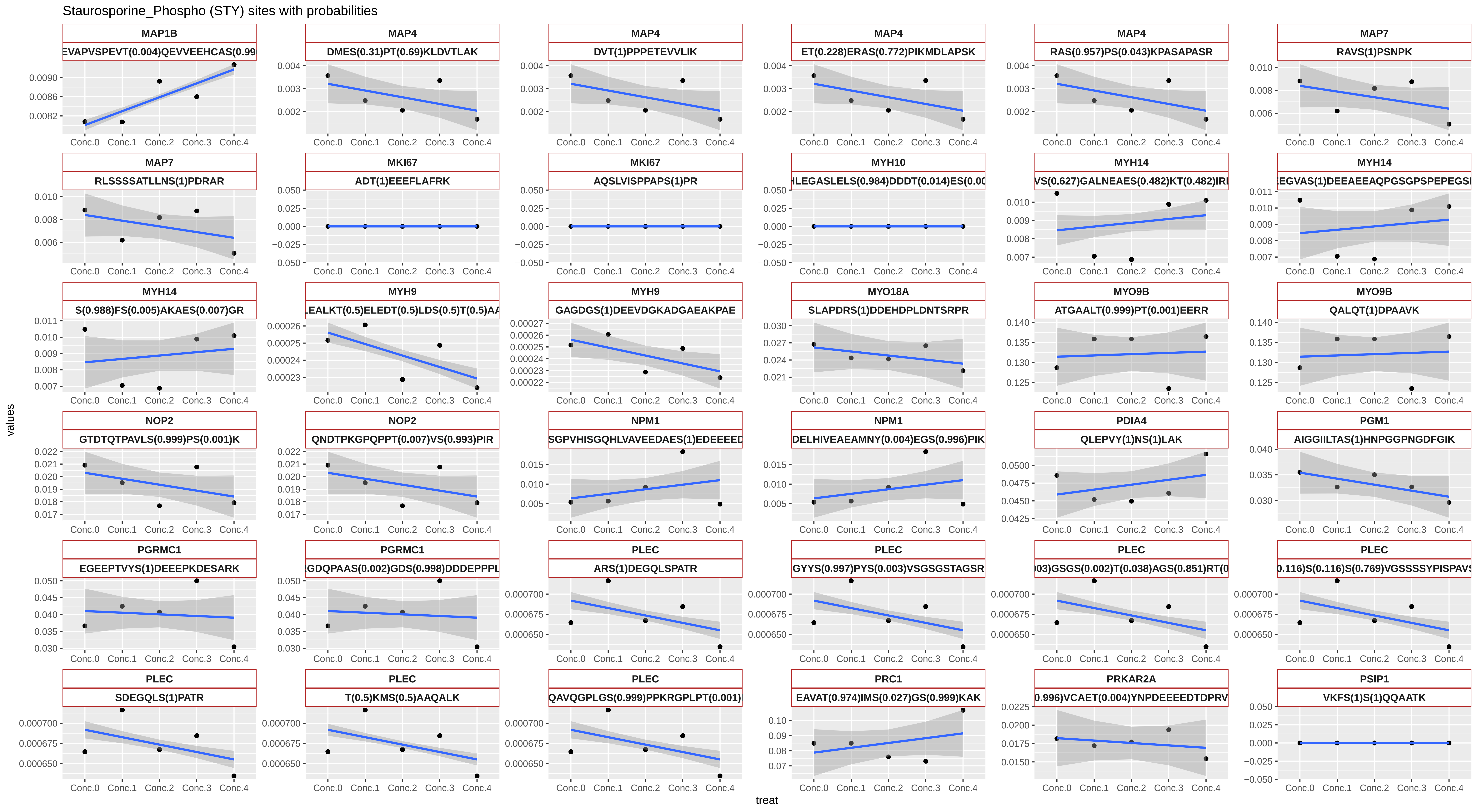

Staurosporine_Phospho (STY) sites with probabilities
MAP1B
EVAPVSPEVT(0.004)QEVVEEHCAS(0.99
MAP4
DMES(0.31)PT(0.69)KLDVTLAK
MAP4
DVT(1)PPPETEVVLIK
MAP4
ET(0.228)ERAS(0.772)PIKMDLAPSK
MAP4
RAS(0.957)PS(0.043)KPASAPASR
MAP7
RAVS(1)PSNPK
MAP7
RLSSSSATLLNS(1)PDRAR
MKI67
ADT(1)EEEFLAFRK
MKI67
AQSLVISPPAPS(1)PR
MYH10
HLEGASLELS(0.984)DDDT(0.014)ES(0.00
MYH14
VS(0.627)GALNEAES(0.482)KT(0.482)IRI
MYH14
EGVAS(1)DEEAEEAQPGSGPSPEPEGSI
MYH14
S(0.988)FS(0.005)AKAES(0.007)GR
MYH9
LEALKT(0.5)ELEDT(0.5)LDS(0.5)T(0.5)AA
MYH9
GAGDGS(1)DEEVDGKADGAEAKPAE
MYO18A
SLAPDRS(1)DDEHDPLDNTSRPR
MYO9B
ATGAALT(0.999)PT(0.001)EERR
MYO9B
QALQT(1)DPAAVK
NOP2
GTDTQTPAVLS(0.999)PS(0.001)K
NOP2
QNDTPKGPQPPT(0.007)VS(0.993)PIR
NPM1
SGPVHISGQHLVAVEEDAES(1)EDEEEED
NPM1
DELHIVEAEAMNY(0.004)EGS(0.996)PIK
PDIA4
QLEPVY(1)NS(1)LAK
PGM1
AIGGIILTAS(1)HNPGGPNGDFGIK
PGRMC1
EGEEPTVYS(1)DEEEPKDESARK
PGRMC1
GDQPAAS(0.002)GDS(0.998)DDDEPPPL
PLEC
ARS(1)DEGQLSPATR
PLEC
GYYS(0.997)PYS(0.003)VSGSGSTAGSR
PLEC
03)GSGS(0.002)T(0.038)AGS(0.851)RT(0
PLEC
0.116)S(0.116)S(0.769)VGSSSSYPISPAVS
PLEC
SDEGQLS(1)PATR
PLEC
T(0.5)KMS(0.5)AAQALK
PLEC
QAVQGPLGS(0.999)PPKRGPLPT(0.001)
PRC1
EAVAT(0.974)IMS(0.027)GS(0.999)KAK
PRKAR2A
0.996)VCAET(0.004)YNPDEEEEDTDPRV
PSIP1
VKFS(1)S(1)QQAATK
Conc.0
Conc.1
Conc.2
Conc.3
Conc.4
values
treat

Staurosporine_Phospho (STY) sites with probabilities

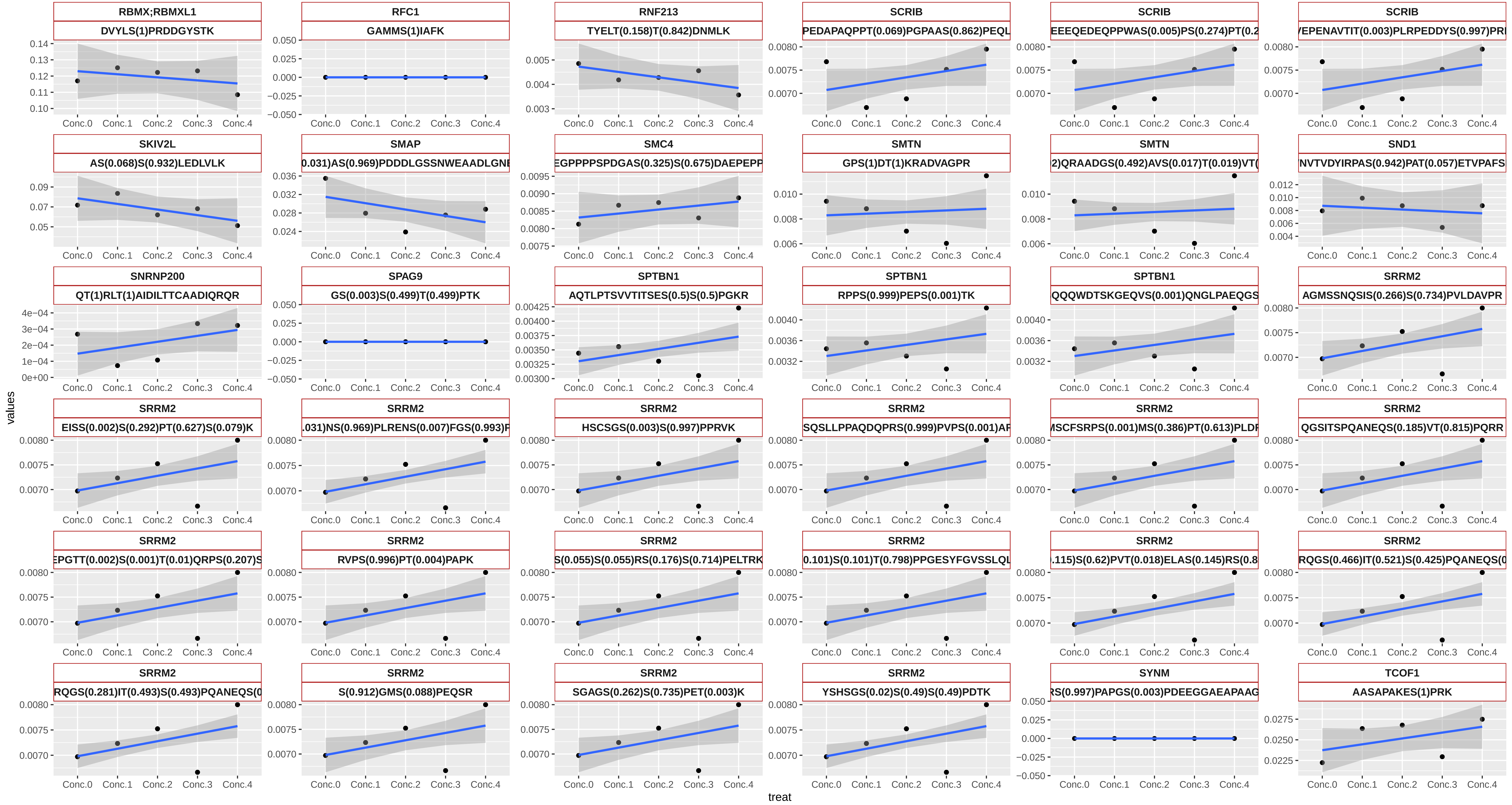

Staurosporine_Phospho (STY) sites with probabilities

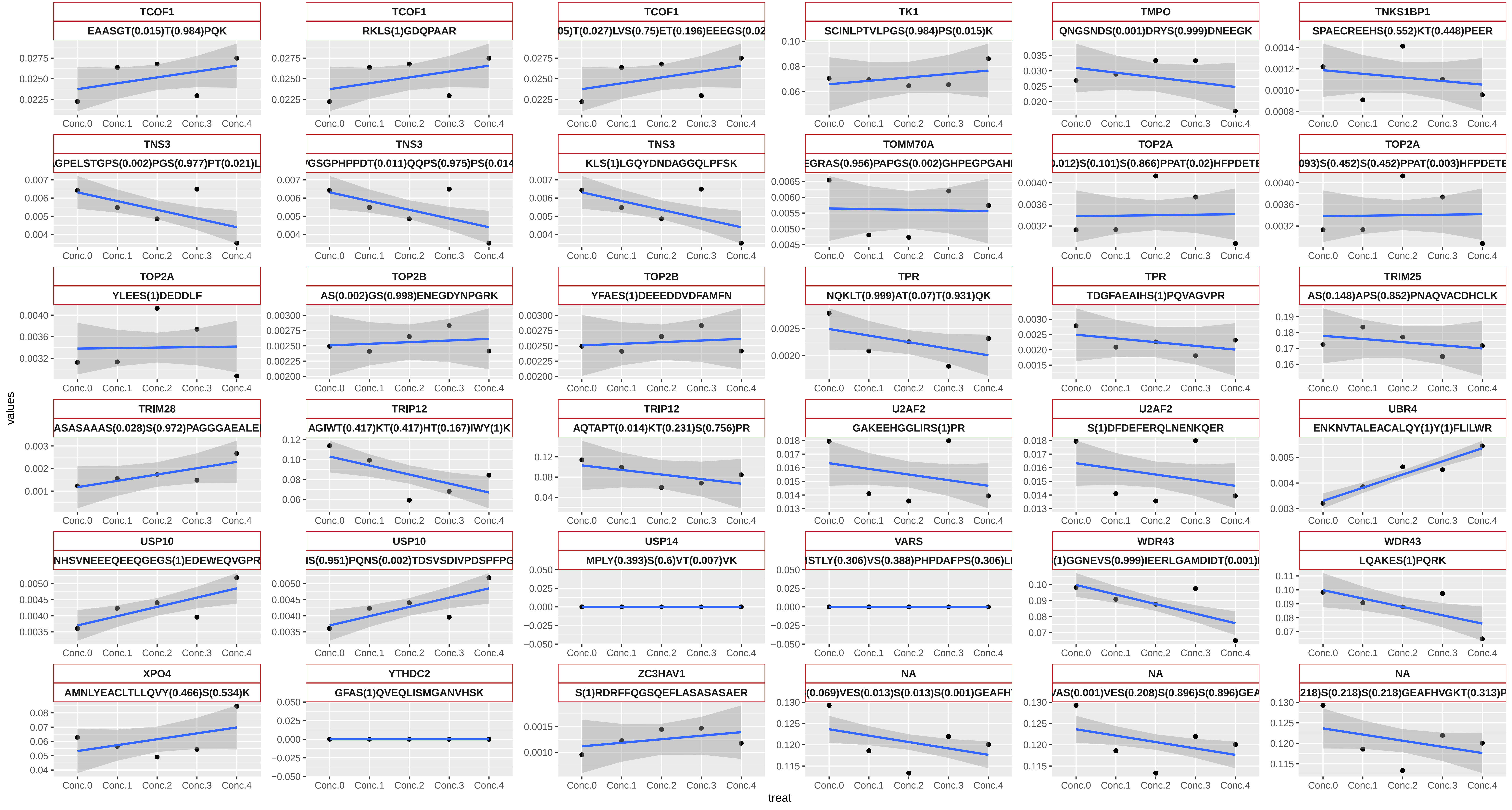

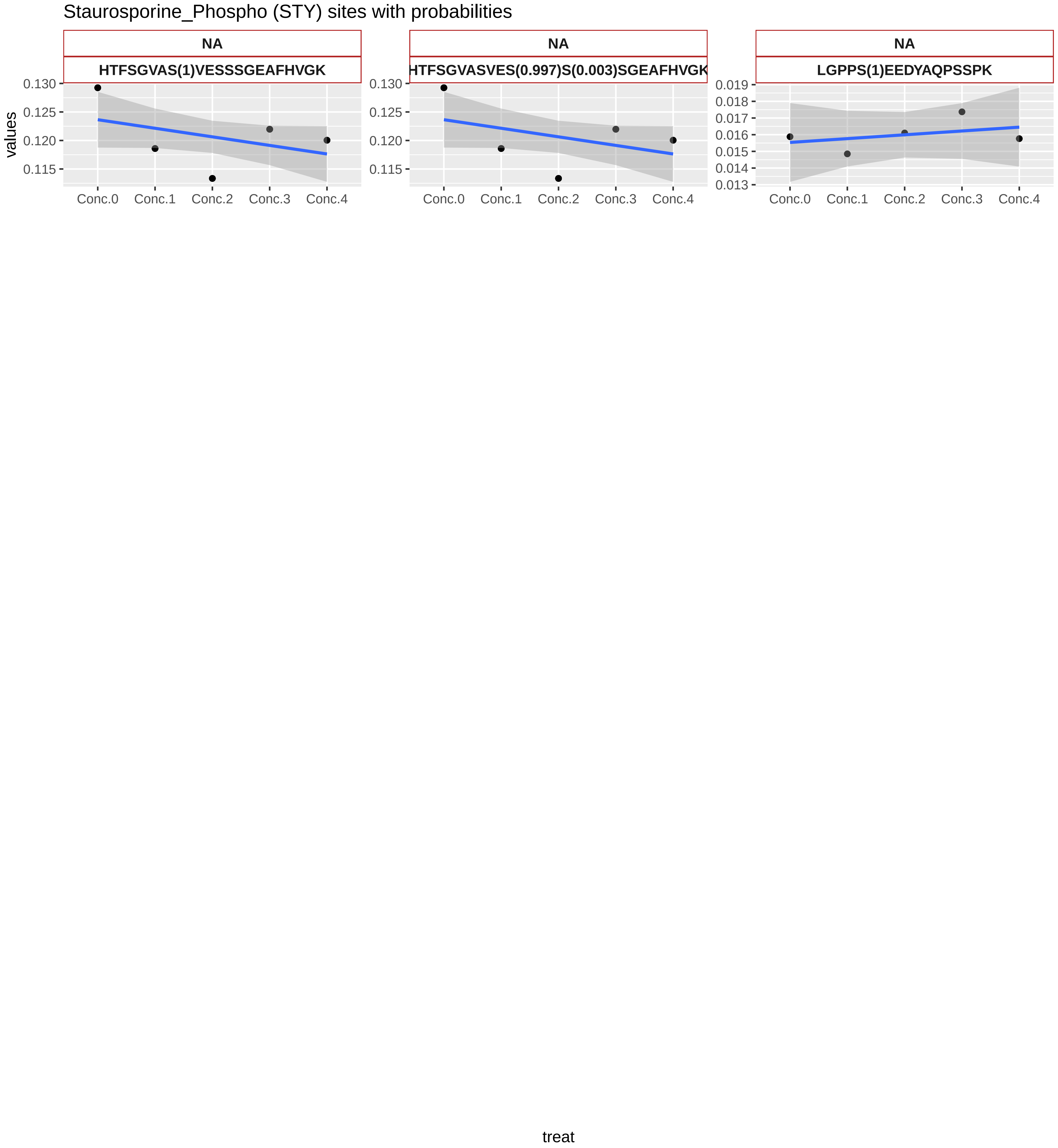

Staurosporine_Phospho (STY) sites with probabilities
NA
HTFSGVAS(1)VESSSGEAFHVGK
NA
HTFSGVASVES(0.997)S(0.003)SGEAFHVGK
NA
LGPPS(1)EEDYAQPSSPK
values
0.130
0.125
0.120
0.115
0.019
0.018
0.017
0.016
0.015
0.014
0.013
Conc.0
Conc.1
Conc.2
Conc.3
Conc.4
treat

Dasatinib_N6−acetyl on Lysine sites with probabilities

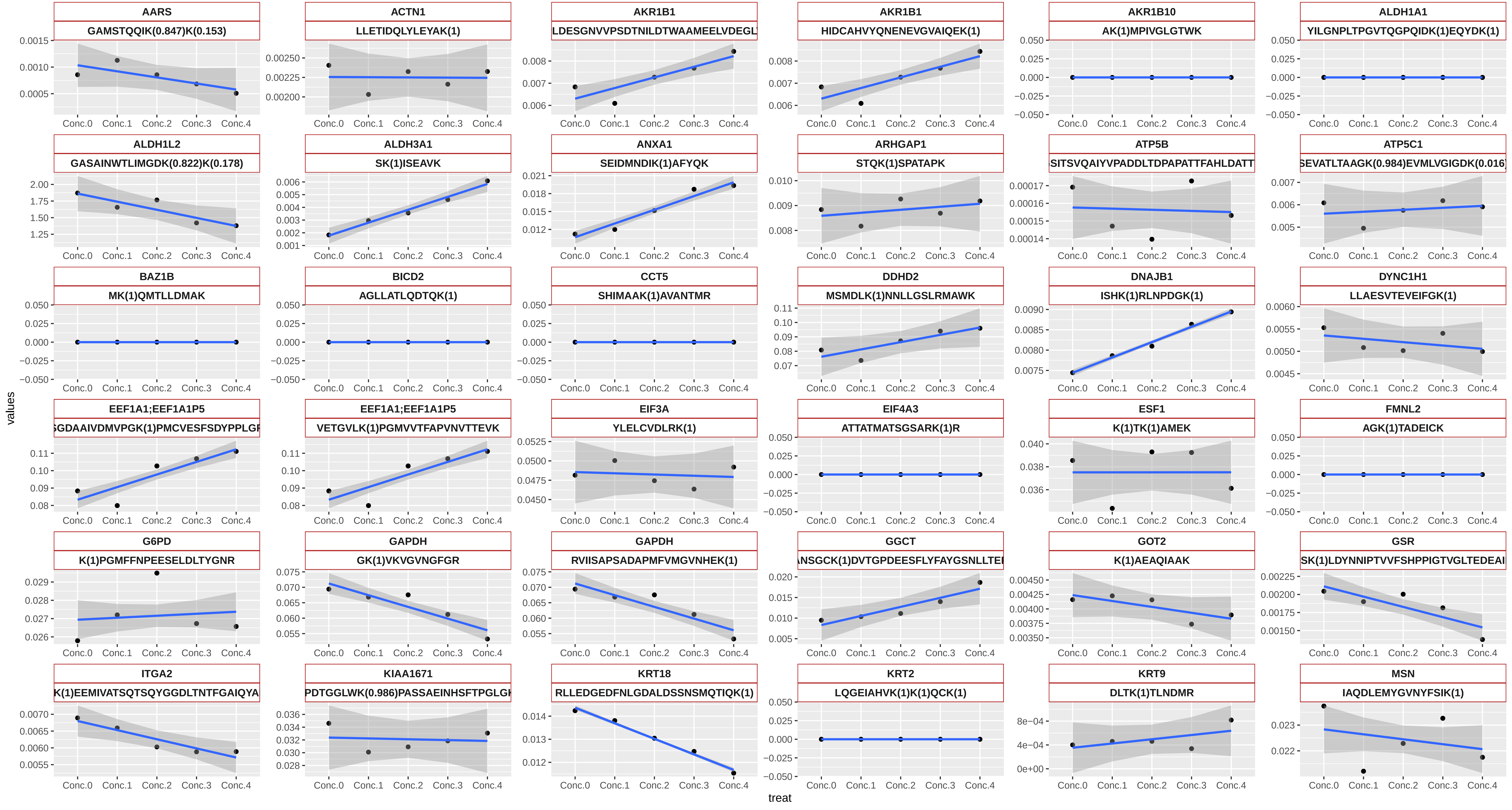

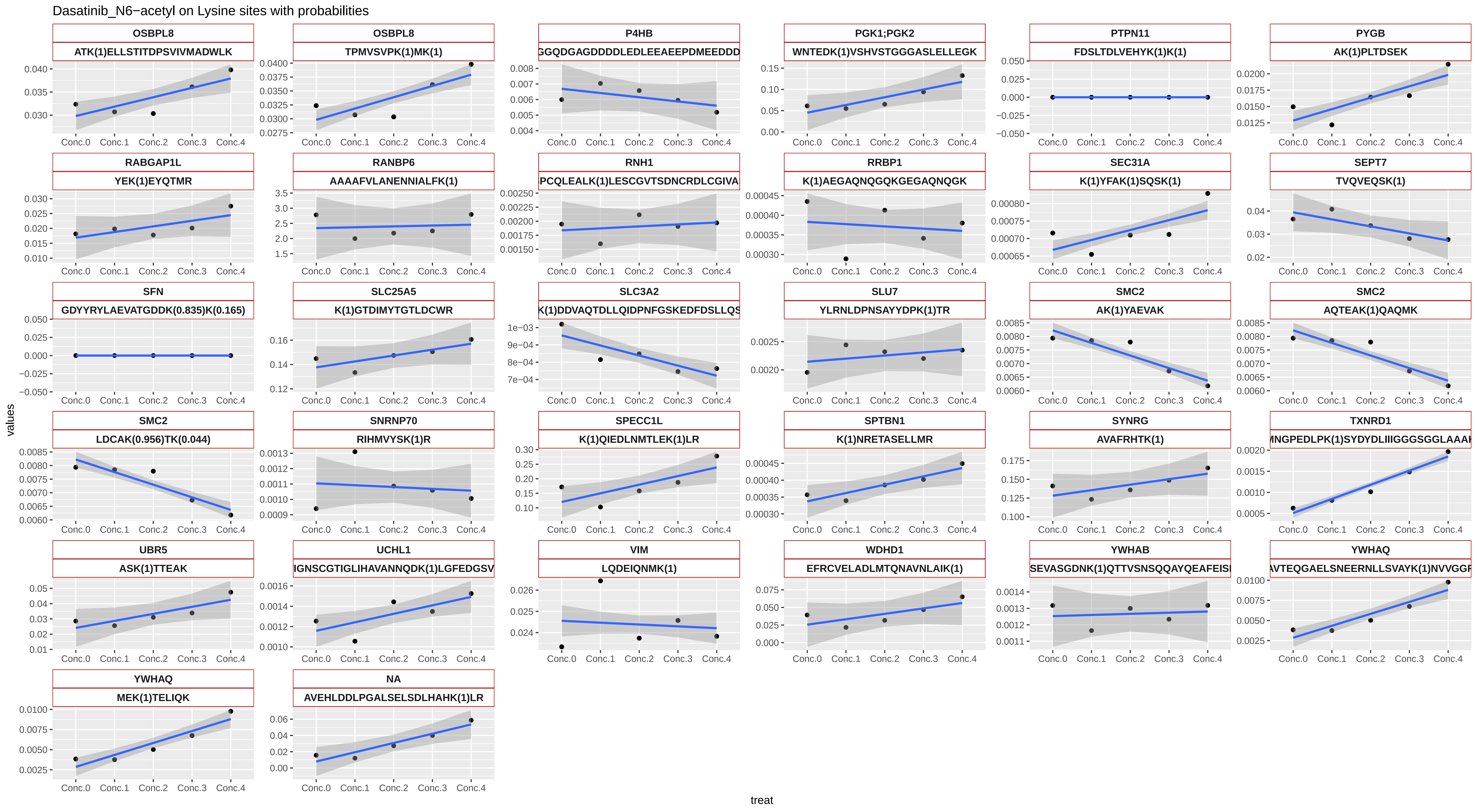
Dasatinib_N6−acetyl on Lysine sites with probabilities
values
treat
OSBPL8
ATK(1)ELLSTITDPSVIVMADWLK
OSBPL8
TPMVSVPK(1)MK(1)
P4HB
GGQDGAGDDDDLEDLEEAEEPDMEEDDD
PGK1;PGK2
WNTEDK(1)VSHVSTGGGASLELLEGK
PTPN11
FDSLTDLVEHYK(1)K(1)
PYGB
AK(1)PLTDSEK
RABGAP1L
YEK(1)EYQTMR
RANBP6
AAAAFVLANENNIALFK(1)
RNH1
PCQLEALK(1)LESCGVTSDNCRDLCGIVA
RRBP1
K(1)AEGAQNQGQKGEGAQNQGK
SEC31A
K(1)YFAK(1)SQSK(1)
SEPT7
TVQVEQSK(1)
SFN
GDYYRYLAEVATGDDK(0.835)K(0.165)
SLC25A5
K(1)GTDIMYTGTLDCWR
SLC3A2
K(1)DDVAQTDLLQIDPNFGSKEDFDSLLQS
SLU7
YLRNLDPNSAYYDPK(1)TR
SMC2
AK(1)YAEVAK
SMC2
AQTEAK(1)QAQMK
SMC2
LDCAK(0.956)TK(0.044)
SNRNP70
RIHMVYSK(1)R
SPECC1L
K(1)QIEDLNMTLEK(1)LR
SPTBN1
K(1)NRETASELLMR
SYNRG
AVAFRHTK(1)
TXNRD1
MNGPEDLPK(1)SYDYDLIIIGGGSGGLAAA
UBR5
ASK(1)TTEAK
UCHL1
IGNSCGTIGLIHAVANNQDK(1)LGFEDGSV
VIM
LQDEIQNMK(1)
WDHD1
EFRCVELADLMTQNAVNLAIK(1)
YWHAB
SEVASGDNK(1)QTTVSNSQQAYQEAFEIS
YWHAQ
AVTEQGAELSNEERNLLSVAYK(1)NVVGG
YWHAQ
MEK(1)TELIQK
NA
AVEHLDDLPGALSELSDLHAHK(1)LR
Conc.0
Conc.1
Conc.2
Conc.3
Conc.4

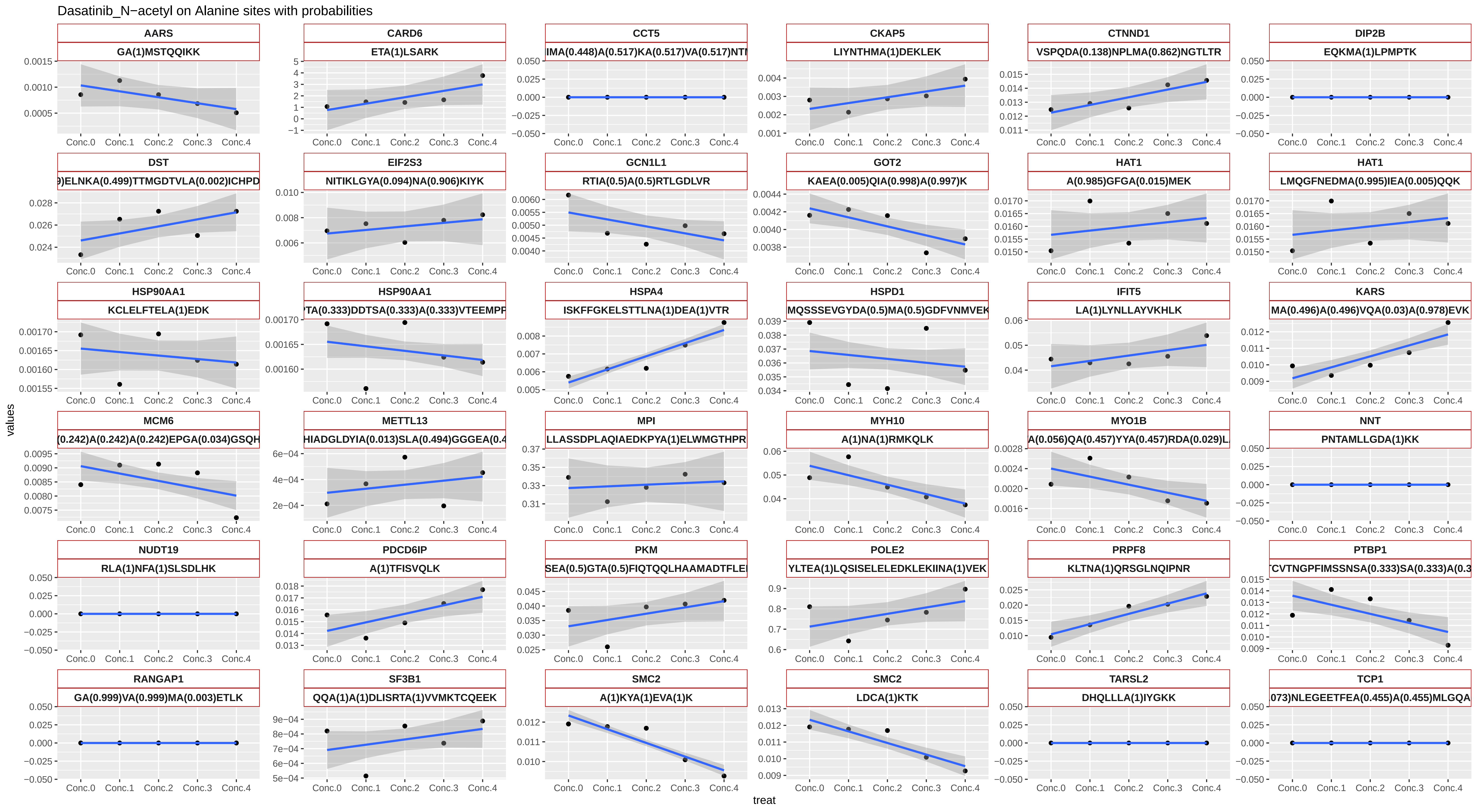

Dasatinib_N−acetyl on Alanine sites with probabilities
AARS
GA(1)MSTQQIKK
CARD6
ETA(1)LSARK
CCT5
IMA(0.448)A(0.517)KA(0.517)VA(0.517)NT
CKAP5
LIYNTHMA(1)DEKLEK
CTNND1
VSPQDA(0.138)NPLMA(0.862)NGTLTR
DIP2B
EQKMA(1)LPMPTK
DST
9)ELNKA(0.499)TTMGDTVLA(0.002)ICHPD
EIF2S3
NITIKLGYA(0.094)NA(0.906)KIYK
GCN1L1
RTIA(0.5)A(0.5)RTLGDLVR
GOT2
KAEA(0.005)QIA(0.998)A(0.997)K
HAT1
A(0.985)GFGA(0.015)MEK
HAT1
LMQGFNEDMA(0.995)IEA(0.005)QQK
HSP90AA1
KCLELFTELA(1)EDK
HSP90AA1
TA(0.333)DDTSA(0.333)A(0.333)VTEEMP
HSPA4
ISKFFGKELSTTLNA(1)DEA(1)VTR
HSPD1
MQSSSEVGYDA(0.5)MA(0.5)GDFVNMVEK
IFIT5
LA(1)LYNLLAYVKHLK
KARS
MA(0.496)A(0.496)VQA(0.03)A(0.978)EVK
MCM6
(0.242)A(0.242)A(0.242)EPGA(0.034)GSQH
METTL13
HIADGLDYIA(0.013)SLA(0.494)GGGEA(0.4
MPI
LLASSDPLAQIAEDKPYA(1)ELWMGTHPR
MYH10
A(1)NA(1)RMKQLK
MYO1B
A(0.056)QA(0.457)YYA(0.457)RDA(0.029)L
NNT
PNTAMLLGDA(1)KK
NUDT19
RLA(1)NFA(1)SLSDLHK
PDCD6IP
A(1)TFISVQLK
PKM
SEA(0.5)GTA(0.5)FIQTQQLHAAMADTFLE
POLE2
YLTEA(1)LQSISELELEDKLEKIINA(1)VEK
PRPF8
KLTNA(1)QRSGLNQIPNR
PTBP1
TCVTNGPFIMSSNSA(0.333)SA(0.333)A(0.3
RANGAP1
GA(0.999)VA(0.999)MA(0.003)ETLK
SF3B1
QQA(1)A(1)DLISRTA(1)VVMKTCQEEK
SMC2
A(1)KYA(1)EVA(1)K
SMC2
LDCA(1)KTK
TARSL2
DHQLLLA(1)IYGKK
TCP1
073)NLEGEETFEA(0.455)A(0.455)MLGQA
Conc.0
Conc.1
Conc.2
Conc.3
Conc.4
values
treat

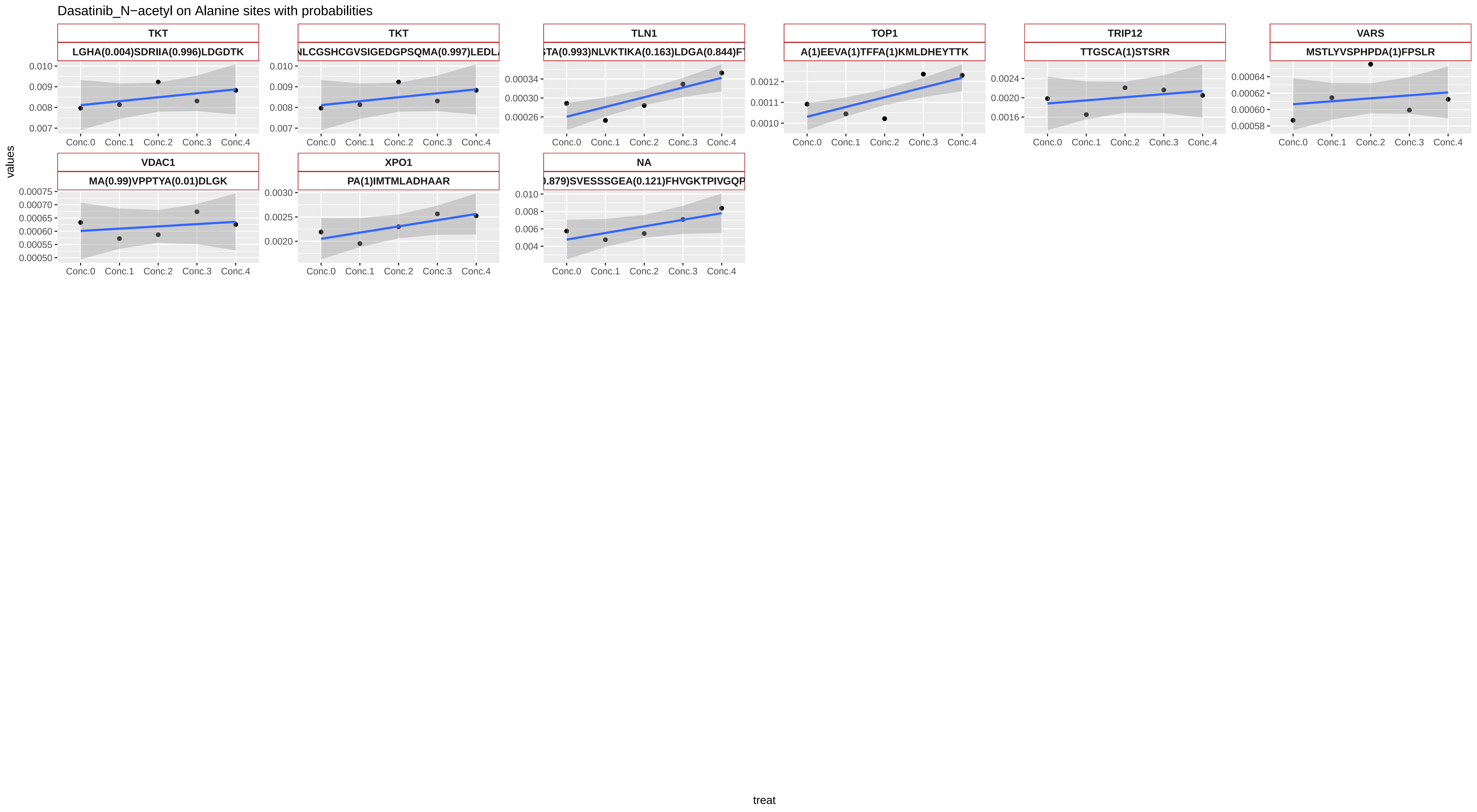

Dasatinib_N−acetyl on Alanine sites with probabilities
TKT
LGHA(0.004)SDRIIA(0.996)LDGDTK
TKT
NLCGSHCGVSIGEDGPSQMA(0.997)LEDL
TLN1
STA(0.993)NLVKTIKA(0.163)LDGA(0.844)F
TOP1
A(1)EEVA(1)TFFA(1)KMLDHEYTTK
TRIP12
TTGSCA(1)STSRR
VARS
MSTLYVSPHPDA(1)FPSLR
VDAC1
MA(0.99)VPPTYA(0.01)DLGK
XPO1
PA(1)IMTMLADHAAR
NA
.879)SVESSSGEA(0.121)FHVGKTPIVGQP
Conc.0
Conc.1
Conc.2
Conc.3
Conc.4
values
treat

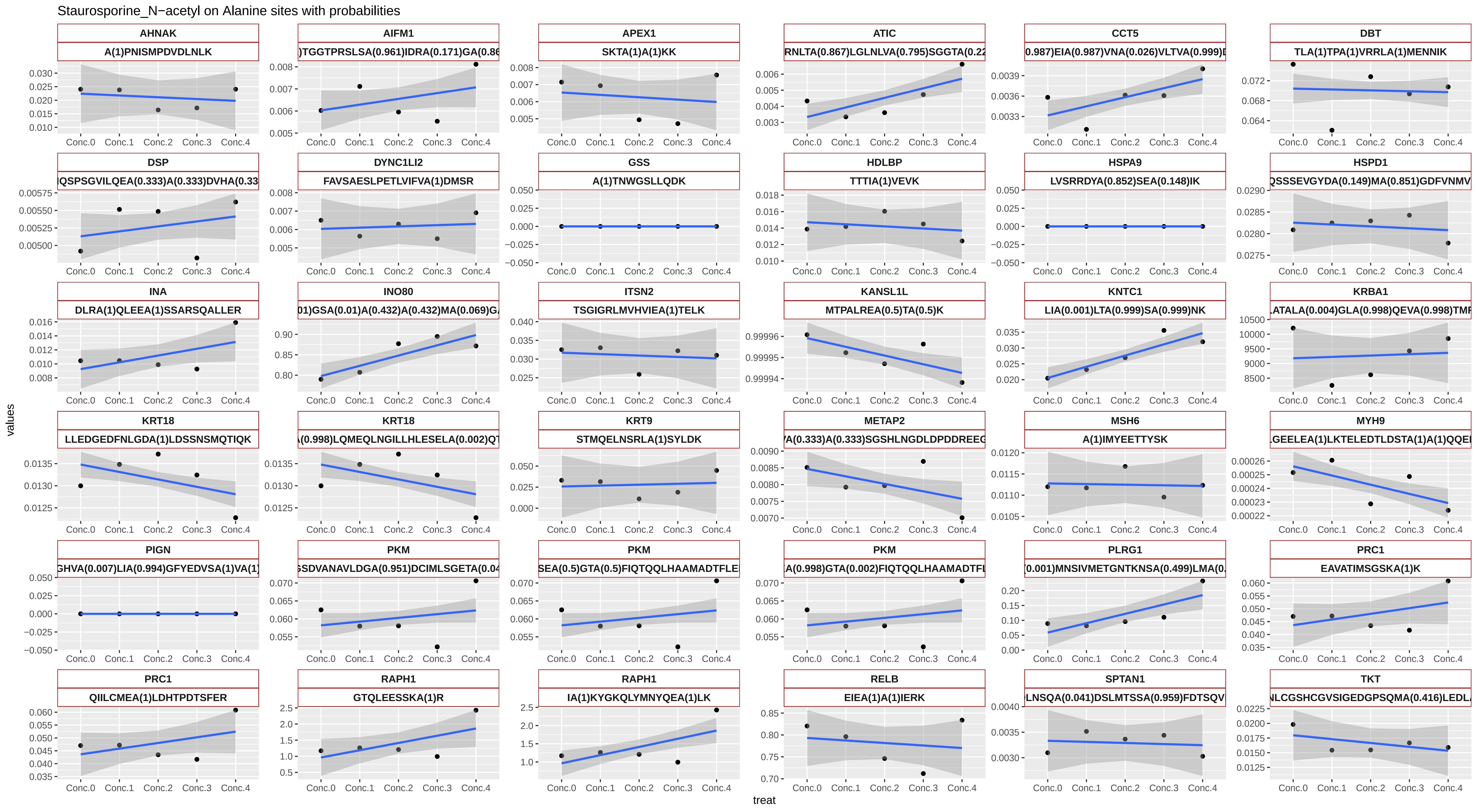

Staurosporine_N−acetyl on Alanine sites with probabilities
AHNAK
A(1)PNISMPDVDLNLK
AIFM1
)TGGTPRSLSA(0.961)IDRA(0.171)GA(0.86
APEX1
SKTA(1)A(1)KK
ATIC
RNLTA(0.867)LGLNLVA(0.795)SGGTA(0.22
CCT5
0.987)EIA(0.987)VNA(0.026)VLTVA(0.999)
DBT
TLA(1)TPA(1)VRRLA(1)MENNIK
DSP
IQSPSGVILQEA(0.333)A(0.333)DVHA(0.33
DYNC1LI2
FAVSAESLPETLVIFVA(1)DMSR
GSS
A(1)TNWGSLLQDK
HDLBP
TTTIA(1)VEVK
HSPA9
LVSRRDYA(0.852)SEA(0.148)IK
HSPD1
QSSSEVGYDA(0.149)MA(0.851)GDFVNMV
INA
DLRA(1)QLEEA(1)SSARSQALLER
INO80
01)GSA(0.01)A(0.432)A(0.432)MA(0.069)G
ITSN2
TSGIGRLMVHVIEA(1)TELK
KANSL1L
MTPALREA(0.5)TA(0.5)K
KNTC1
LIA(0.001)LTA(0.999)SA(0.999)NK
KRBA1
LATALA(0.004)GLA(0.998)QEVA(0.998)TMF
KRT18
LLEDGEDFNLGDA(1)LDSSNSMQTIQK
KRT18
A(0.998)LQMEQLNGILLHLESELA(0.002)QT
KRT9
STMQELNSRLA(1)SYLDK
METAP2
VA(0.333)A(0.333)SGSHLNGDLDPDDREEG
MSH6
A(1)IMYEETTYSK
MYH9
LGEELEA(1)LKTELEDTLDSTA(1)A(1)QQE
PIGN
GHVA(0.007)LIA(0.994)GFYEDVSA(1)VA(1)
PKM
SGSDVANAVLDGA(0.951)DCIMLSGETA(0.04
PKM
SEA(0.5)GTA(0.5)FIQTQQLHAAMADTFLE
PKM
A(0.998)GTA(0.002)FIQTQQLHAAMADTFL
PLRG1
(0.001)MNSIVMETGNTKNSA(0.499)LMA(0.
PRC1
EAVATIMSGSKA(1)K
PRC1
QIILCMEA(1)LDHTPDTSFER
RAPH1
GTQLEESSKA(1)R
RAPH1
IA(1)KYGKQLYMNYQEA(1)LK
RELB
EIEA(1)A(1)IERK
SPTAN1
LNSQA(0.041)DSLMTSSA(0.959)FDTSQV
TKT
NLCGSHCGVSIGEDGPSQMA(0.416)LEDL
Conc.0
Conc.1
Conc.2
Conc.3
Conc.4
values
treat

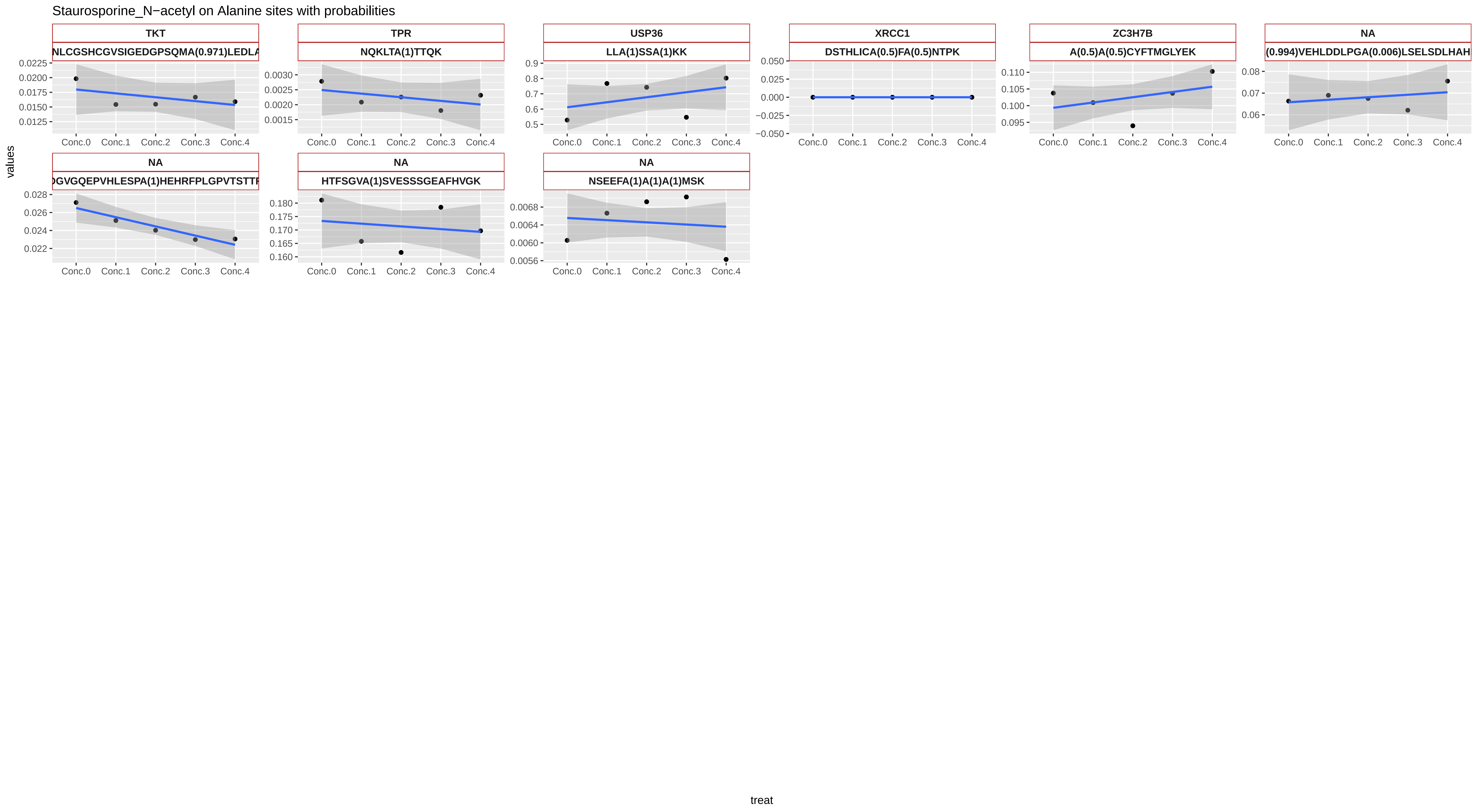
Staurosporine_N−acetyl on Alanine sites with probabilities
TKT
NLCGSHCGVSIGEDGPSQMA(0.971)LEDLA
TPR
NQKLTA(1)TTQK
USP36
LLA(1)SSA(1)KK
XRCC1
DSTHLICA(0.5)FA(0.5)NTPK
ZC3H7B
A(0.5)A(0.5)CYFTMGLYEK
NA
(0.994)VEHLDDLPGA(0.006)LSELSDLHAH
NA
DGVGQEPVHLESPA(1)HEHRFPLGPVTSTTR
NA
HTFSGVA(1)SVESSSGEAFHVGK
NA
NSEEFA(1)A(1)A(1)MSK
Conc.0
Conc.1
Conc.2
Conc.3
Conc.4
values
treat

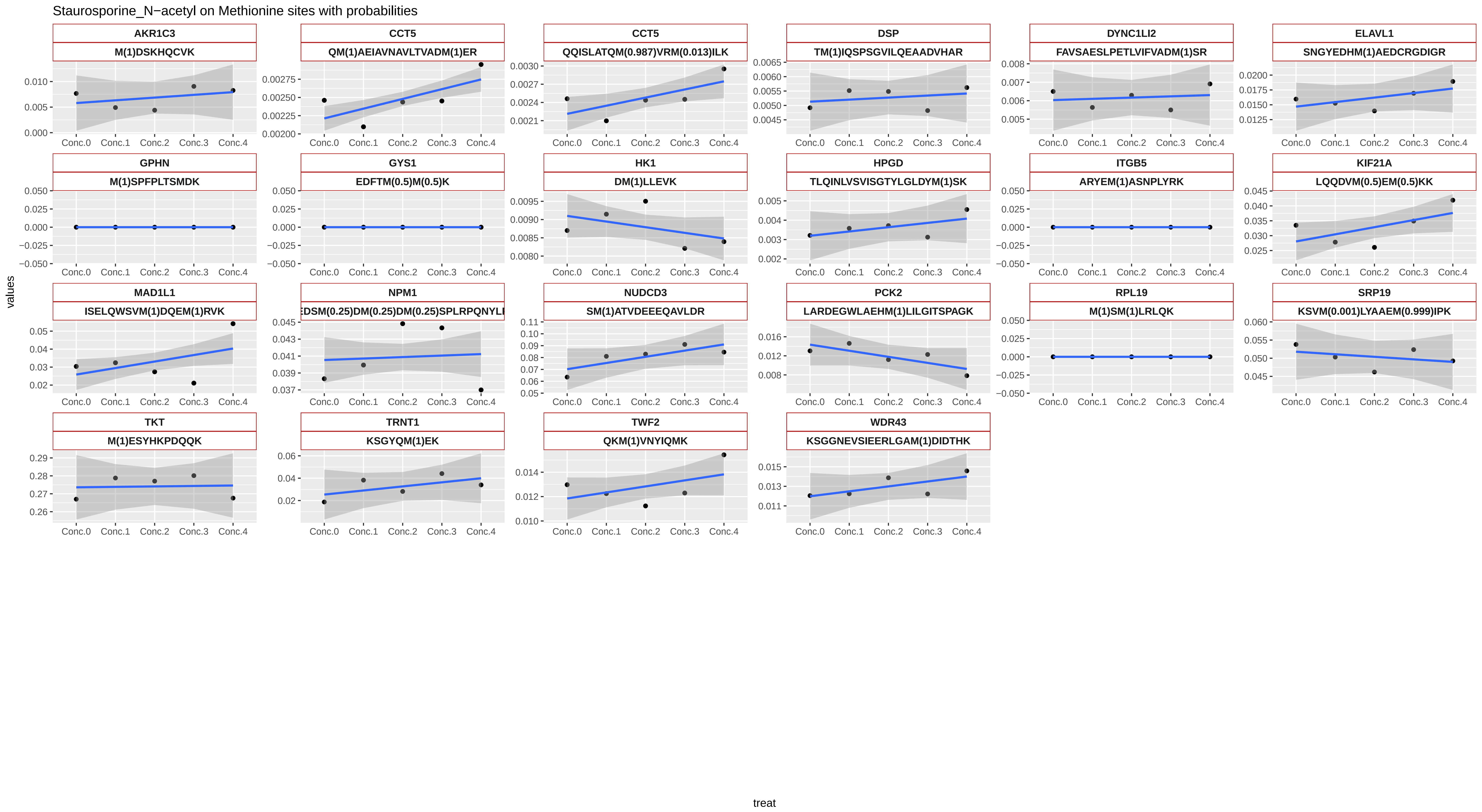
Staurosporine_N−acetyl on Methionine sites with probabilities
AKR1C3
M(1)DSKHQCVK
CCT5
QM(1)AEIAVNAVLTVADM(1)ER
CCT5
QQISLATQM(0.987)VRM(0.013)ILK
DSP
TM(1)IQSPSGVILQEAADVHAR
DYNC1LI2
FAVSAESLPETLVIFVADM(1)SR
ELAVL1
SNGYEDHM(1)AEDCRGDIGR
GPHN
M(1)SPFPLTSMDK
GYS1
EDFTM(0.5)M(0.5)K
HK1
DM(1)LLEVK
HPGD
TLQINLVSVISGTYLGLDYM(1)SK
ITGB5
ARYEM(1)ASNPLYRK
KIF21A
LQQDVM(0.5)EM(0.5)KK
MAD1L1
ISELQWSVM(1)DQEM(1)RVK
NPM1
EDSM(0.25)DM(0.25)DM(0.25)SPLRPQNYLI
NUDCD3
SM(1)ATVDEEEQAVLDR
PCK2
LARDEGWLAEHM(1)LILGITSPAGK
RPL19
M(1)SM(1)LRLQK
SRP19
KSVM(0.001)LYAAEM(0.999)IPK
TKT
M(1)ESYHKPDQQK
TRNT1
KSGYQM(1)EK
TWF2
QKM(1)VNYIQMK
WDR43
KSGGNEVSIEERLGAM(1)DIDTHK
Conc.0
Conc.1
Conc.2
Conc.3
Conc.4
values
treat
